# Processing macromolecular diffuse scattering data

**DOI:** 10.1101/2023.06.04.543637

**Authors:** Steve P. Meisburger, Nozomi Ando

## Abstract

Diffuse scattering is a powerful technique to study disorder and dynamics of macromolecules at atomic resolution. Although diffuse scattering is always present in diffraction images from macromolecular crystals, the signal is weak compared with Bragg peaks and background, making it a challenge to visualize and measure accurately. Recently, this challenge has been addressed using the reciprocal space mapping technique, which leverages ideal properties of modern X-ray detectors to reconstruct the complete three-dimensional volume of continuous diffraction from diffraction images of a crystal (or crystals) in many different orientations. This chapter will review recent progress in reciprocal space mapping with a particular focus on the strategy implemented in the *mdx-lib* and *mdx2* software packages. The chapter concludes with an introductory data processing tutorial using Python packages *DIALS, NeXpy*, and *mdx2*.

## 1. Introduction

The fact that proteins and nucleic acids form crystals that diffract to (near) atomic resolution has had a profound impact on the biological sciences. X-ray diffraction data from macromolecular crystals is responsible for over 170,000 structures in the Protein Data Bank, the majority at 2.5 Å resolution or better. Although the X-ray scattering from individual molecules is weak, when they are aligned in a crystal, the diffraction interferes constructively in certain directions, causing sharp Bragg peaks to appear in diffraction images. The Bragg peak intensities depend on the repeating pattern of electron density in the crystal, and thus they are used for structure determination. However, not all X-rays scattered by a crystal go into Bragg peaks. In fact, for most protein crystals, the majority of the diffracted X-rays are found elsewhere, in a continuous diffuse pattern (Clarage & Phillips Jr, 1997). Bragg diffraction and diffuse scattering are complementary: the former depends on regularity, the latter on deviations from the average. Intriguingly, the pattern of diffuse scattering encodes the statistical correlations of electron density fluctuations (Meisburger et al., 2017). Such information is urgently needed in the modern era of structural biology, which seeks to explain function in terms of conformational dynamics, excited states, and ensembles (Ourmazd et al., 2022). This need has motivated a renewed effort to develop methods to accurately measure diffuse scattering for studies of correlated motion (Xu et al., 2021).

Recently, we reported detailed maps of diffuse scattering from lysozyme in triclinic, orthorhombic, and tetragonal space groups (Meisburger et al., 2020, 2023). These high quality crystals had sufficiently low mosaicity at room temperature to finely sample the diffuse scattering close to the Bragg peaks. The lysozyme datasets contain two features: a weak, slowly varying cloudy pattern, and intense halos of scattering around the Bragg peaks. Similar halo features have been observed in all diffuse maps reported to date with sufficient accuracy and precision to resolve them (De Klijn et al., 2019; Glover et al., 1991; Moore, 2009; Peck et al., 2018; Wall, Clarage, et al., 1997), suggesting that halos are a universal feature of protein crystal diffraction. In lysozyme, the intensities and shapes of the halo features could be accounted for by modeling the vibrations of the crystal lattice. By measuring and accounting for the halos, the weaker residual scattering could be analyzed to fit models of internal motion (Meisburger et al., 2020).

The presence of halos in protein diffuse scattering has important implications for data processing. First, to resolve the halos accurately, the three dimensional maps must be finely sampled. Lattice disorder models applicable to protein crystals predict halo shapes characterized by a sharp decay away from the Bragg peak on a scale much finer than the reciprocal lattice spacing. Second, the cloudy signal of interest is weak and partly obscured by the more intense halos, and therefore experimental artifacts that produce modulations in background scattering must be minimized experimentally or otherwise corrected in software. In this chapter, we first give a brief history of continuous diffraction measurements from protein crystals and review the diffraction theory relevant to data processing. We describe how data processing challenges are addressed computationally with a particular emphasis on the approach taken in *mdx-lib* and *mdx2*. Finally, we present an introductory tutorial where *mdx2* is used to reconstruct the three dimensional map from a cubic insulin dataset collected at ambient temperature.

### 1.1. Brief history of macromolecular diffuse scattering

Macromolecular diffuse scattering has a long history (Meisburger et al., 2017). The modern approach of quantitative reciprocal space mapping (i.e., the process of not only mapping the detector coordinates to reciprocal space but also correcting intensities to accurately represent the crystal’s elastic scattering) is rooted in a series of visionary studies by Donald Caspar and colleagues, beginning with their 1988 letter to Nature reporting the diffuse scattering from insulin (Caspar et al., 1988). The diffraction pattern from a cubic insulin crystal was recorded on X-ray film in a 40-h exposure (Bragg peaks were overexposed), digitized, and processed to correct for background, polarization, and the Compton and water scattering contributions (Figure 1A). The processing revealed a diffuse pattern with two components: halos centered around the Bragg peaks, and cloudy “variational” scattering background. The halos were interpreted as displacement disorder of the lattice with correlations that decay exponentially with distance between unit cells. The variational scattering was explained using a similar function, but with correlated displacements between atoms in the protein. Importantly, Caspar and colleagues demonstrated both that accurate measurements of diffuse scattering are feasible, and the technique’s potential to reveal protein dynamics.

**Figure 1.**
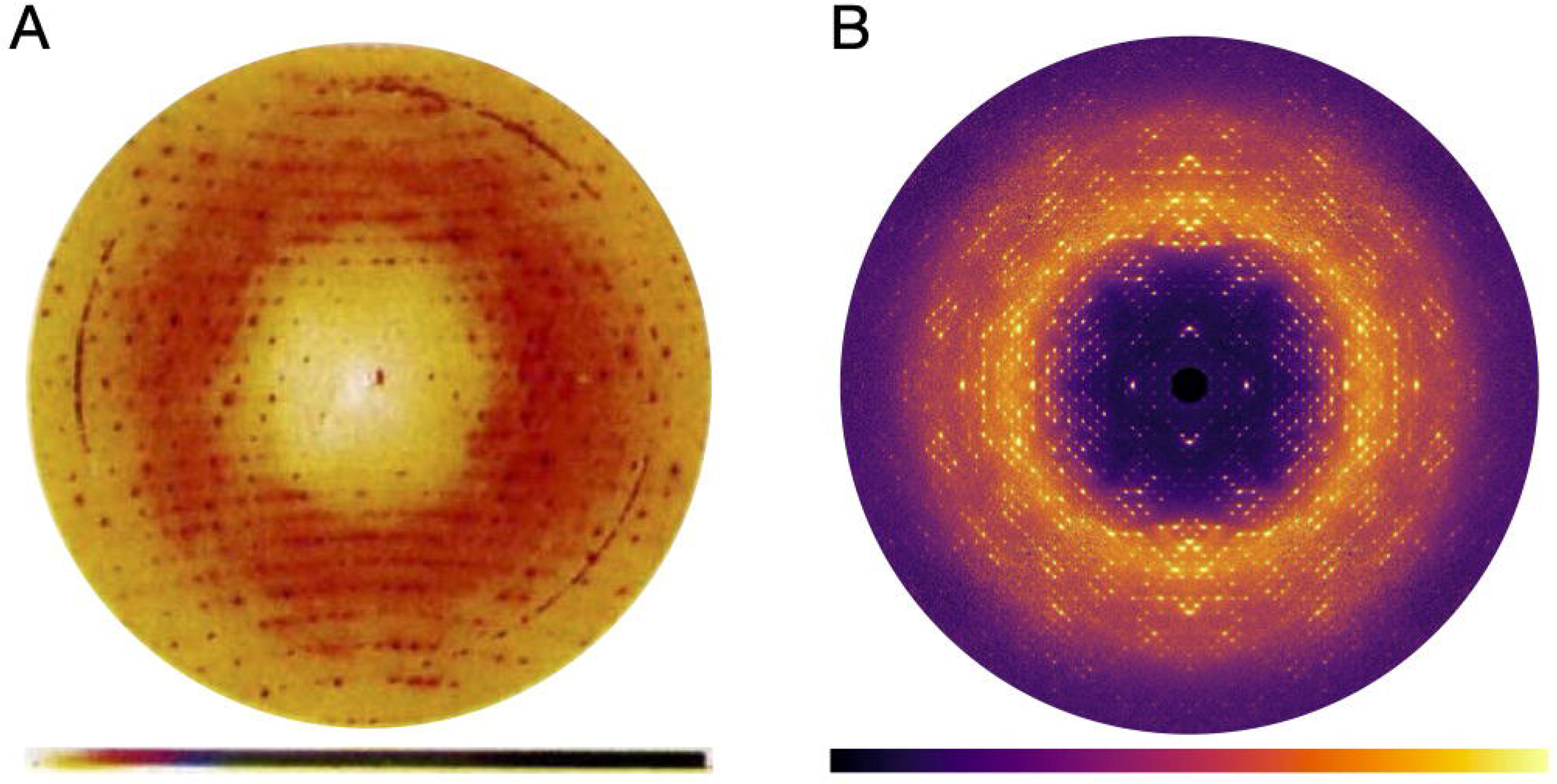
Reciprocal space maps of cubic insulin. (A) A diffraction image recorded on X-ray film, digitized, and processed to remove background and polarization effects. A cloudy pattern and halos surrounding the Bragg peaks are superimposed on the largely isotropic diffuse scattering. Ring features are background scattering from the aluminum windows. Reproduced with permission from (Caspar et al., 1988). (B) Central section (hk0) through the diffuse scattering map reconstructed using *mdx2* from fine-sliced rotation data collected using a photon-counting detector (see tutorial in Section 3).

Caspar and his contemporaries analyzed two dimensional diffraction patterns. Mathematically, such patterns sample a curved surface (the Ewald sphere) through a three-dimensional reciprocal space function aligned with the crystal. The next methodological breakthrough was to reconstruct the three dimensional function from many diffraction patterns collected with the crystal at different orientations. The first such method was developed by Michael Wall in the lab of Sol Gruner, and it used a modified charge-coupled device (CCD) detector to record the diffuse signal without interference from over-exposed Bragg peaks. Computer memory limitations at the time made it infeasible to reconstruct the complete map at a sampling rate sufficient to resolve the halos (Wall, Clarage, et al., 1997), however coarse maps were generated for crystals of staphylococcal nuclease (Wall, Ealick, et al., 1997), and later, the halos of a calmodulin crystal were resolved in a limited region (Wall, Clarage, et al., 1997). Three-dimensional mapping transformed the way experiment was compared with simulation and with data from other research groups; diffuse scattering became a quantitative and reproducible measurement (Wall, 1996; Wall, Ealick, et al., 1997), and statistics were developed to judge model quality analogous to R-factors in crystallography (Wall, Ealick, et al., 1997).

Modern approaches to reciprocal space mapping benefit from the ideal properties of hybrid photon-counting (HPC) detectors. Unlike CCDs and X-ray film from prior eras, the dynamic range of modern detectors is sufficient to record both Bragg peaks and diffuse scattering in the same image. Because of single-photon sensitivity, long exposures are not needed to overcome the intrinsic detector noise, and the diffuse signal can be accumulated over many observations. For instance, the reciprocal space maps of lysozyme mentioned in the previous section were reconstructed from fine-sliced datasets of 0.1 degrees per frame where most pixels recorded zero or one photon.

The introduction of intense X-ray free electron laser (XFEL) sources in the last decade has spurred interest in dynamic crystallography at non-cryogenic conditions (Shoemaker & Ando, 2018). Bright X-ray pulses from XFELs and fourth generation synchrotrons are sufficient to destroy the crystal in a single shot, requiring many crystals to obtain a complete dataset. At XFELs in particular, single-shot measurements of very small crystals are possible because the pulse duration is faster than the radiation damage process. XFELs have also proven useful for measuring very weak, non-Bragg scattering signals. For instance, interference fringes representing the crystal’s shape transform have been visualized between the Bragg peaks of Photosystem I nanocrystals (Chapman et al., 2011). From larger crystals of Photosystem II, diffuse scattering beyond the limits of the Bragg diffraction resolution were observed in XFEL diffraction patterns and the diffuse scattering was used to improve the resolution of the electron density through iterative phasing techniques (Ayyer et al., 2016; Morgan et al., 2019).

### 1.2 Data Processing Software

Software is a key component of the reciprocal space mapping technique. A variety of software tools have been developed (Table 1). Although each is specialized for particular applications or mode of data collection, a common workflow emerges. First, the Bragg peak positions are analyzed using an indexing program such as *XDS* (Kabsch, 2010) or *DIALS* (Winter et al., 2022) to determine the orientation of the crystal during each X-ray exposure. Using this information, the pixels in each image are assigned reciprocal space coordinates. The data are further corrected for geometric effects such as solid angle and polarization. Finally, the corrected data are merged on a regular, three-dimensional grid, yielding a reciprocal space map.

**Table 1.**
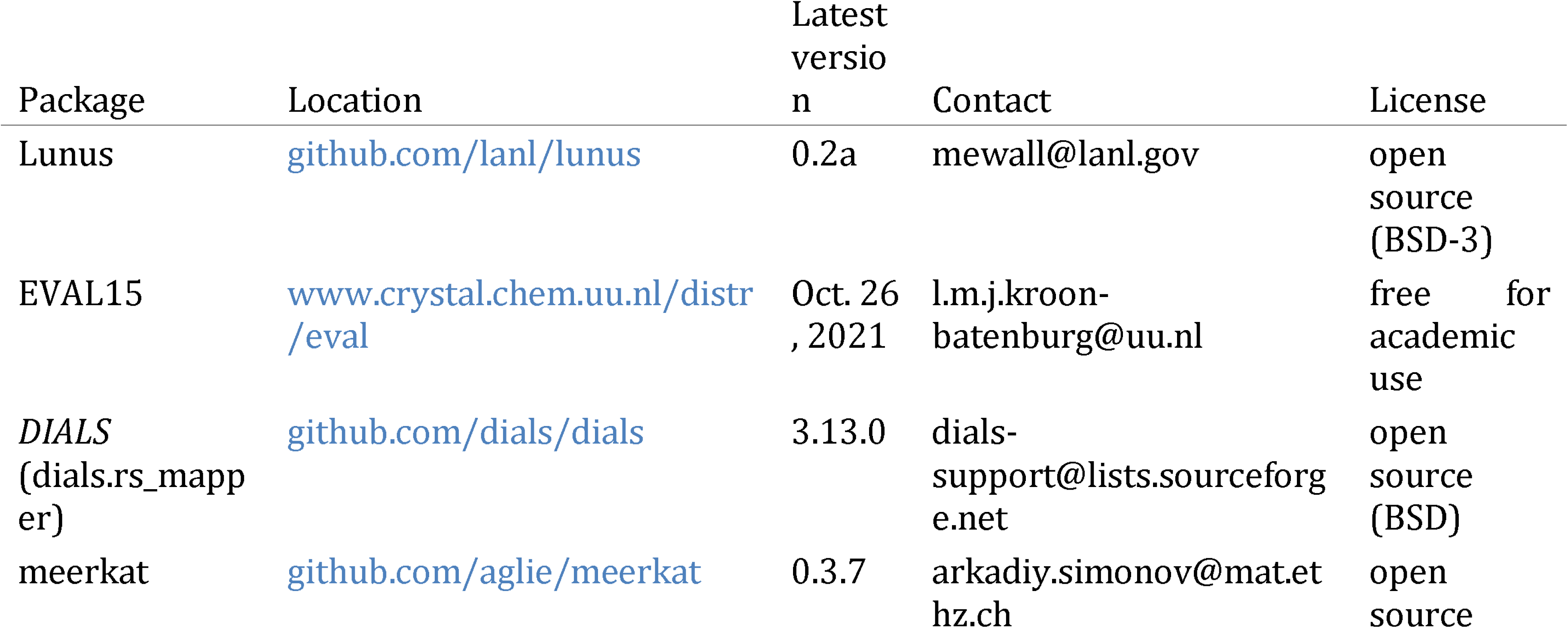
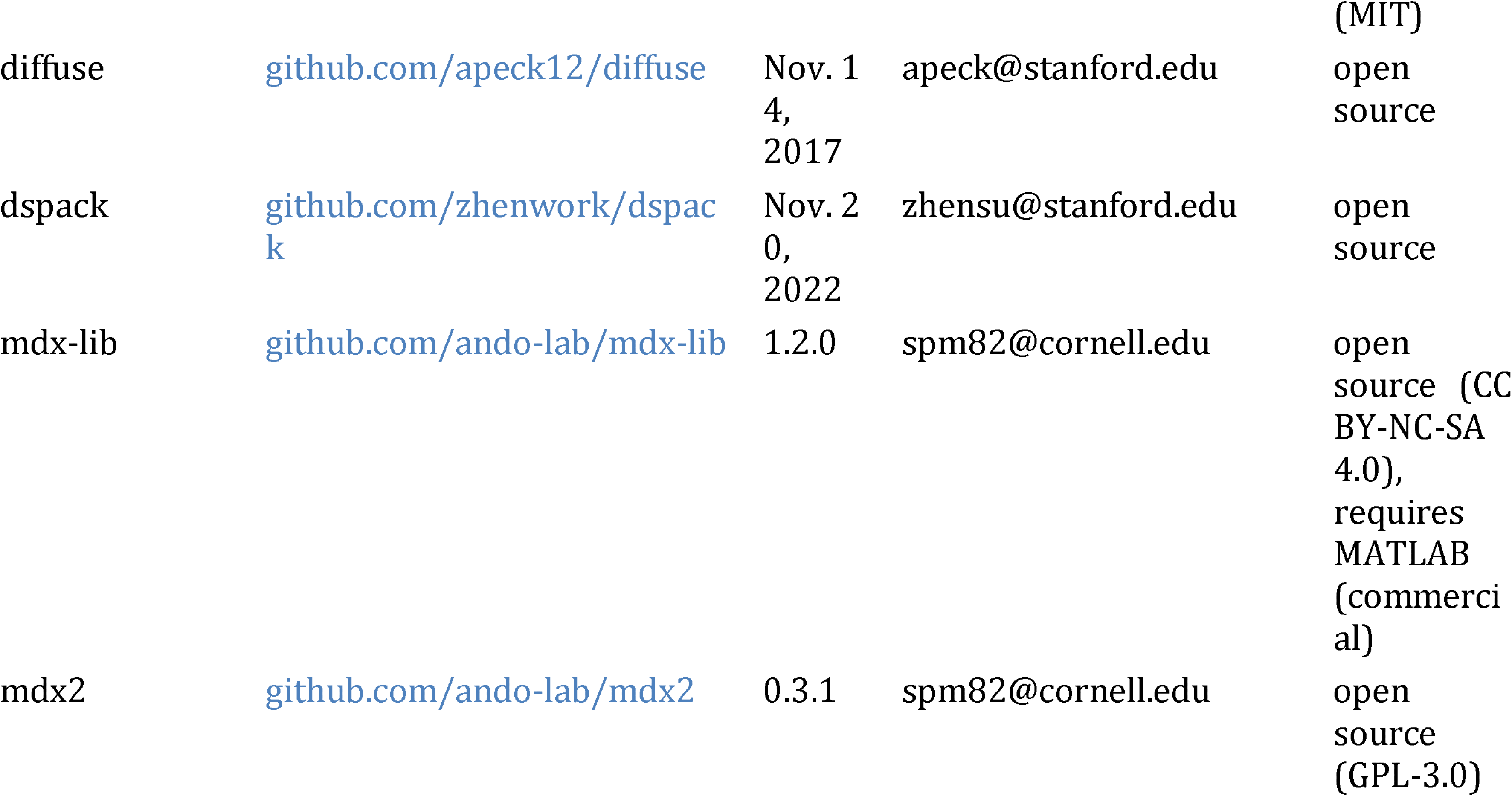
Software for reciprocal space reconstruction.

Despite following similar workflows, current software tools differ in the types of experiments they can analyze, and the details of how experimental artifacts are treated. Thus, when choosing software it is important to consider the application. When reciprocal space mapping is used to diagnose or otherwise visualize pathological crystal packing, accuracy is less important than ease of use. One particularly convenient program is *dials*.*rs_mapper*, which generates 3D maps from indexed diffraction data (Winter et al., 2022). The output format for the 3D map is compatible with software developed for viewing electron density. The *EVAL* suite of programs (De Klijn et al., 2019; Schreurs et al., 2010) should be used when greater detail is needed, for instance to visualize sharp streaks or satellite reflections from modulated structures (Neviani et al., 2022). *EVAL* implements pixel splitting to produce densely sampled maps, and it includes an array of additional tools for analyzing non-standard diffraction data. Several software packages deal specifically with macromolecular diffuse scattering, and each has unique features. *Lunus* (Van Benschoten et al., 2016; Wall, 2009) has been optimized for high-performance computing architectures. *Diffuse* implements a novel scaling and background subtraction algorithm based on principal component analysis (Peck et al., 2018). *Dspack* has a streamlined command-line interface (Su et al., 2021). *Mdx-lib* is particularly suited to high-redundancy data, and it includes methods for background subtraction, scaling model refinement, and error propagation (Meisburger et al., 2020). Finally, diffuse scattering software designed for non-biological work is also available. In particular, *meerkat* (Simonov et al., 2020) has many similarities with the biological software tools described above.

Accessibility is also a consideration when choosing software. Most diffuse scattering projects are open source with permissive licenses (see Table 1). However, the level of effort required to use the software effectively on a new project is typically high. The user base for each diffuse scattering program is currently small, in most cases limited to the research group that developed it. Thus, there has not been a need to build general-purpose tools, streamlined interfaces, tutorials, or thorough documentation. For example, analyzing a new dataset using *mdx-lib* requires a high level of expertise in MATLAB to understand and modify example scripts. It is also important to note that many packages are in active development, and collaboration with the authors of the code may be required to implement necessary features, such as detector image formats.

As biological diffuse scattering transitions from a proof-of-concept experiment to a widely available technique, software tools are needed that are rigorous but user-friendly. Since experimental and algorithic techniques are likely to evolve, the software should also be easy to extend and modify. Moreover, the software should be developed collaboratively and maintained by the community. The ideal software does not yet exist. Motivated by this need, we have developed a successor to *mdx-lib* called *mdx2*, which we describe for the first time in this chapter (Figure 2). *Mdx2* re-implements the data processing workflow of *mdx-lib* in python, but it does so without reinventing crystallographic computing (as *mdx-lib* did). Underneath, *mdx2* is lightweight, relying on *dxtbx* (Parkhurst et al., 2014) for data import and *cctbx* (Grosse-Kunstleve et al., 2002) for crystallographic computing. Since *DIALS* uses the same packages (Winter et al., 2022), *mdx2* and *DIALS* functions are interoperable. *Mdx2* includes a command-line interface (also inspired by *DIALS*), so familiarity with python is not required. For more flexible workflows, *mdx2* can be imported as a python package and run for instance in a Jupyter notebook. Finally, we have chosen to store all intermediate data files in *NeXus* format (Könnecke et al., 2015) to make them interoperable with other scientific software, especially the data visualization program *NeXpy* (The NeXpy development team, 2023).

**Figure 2.**
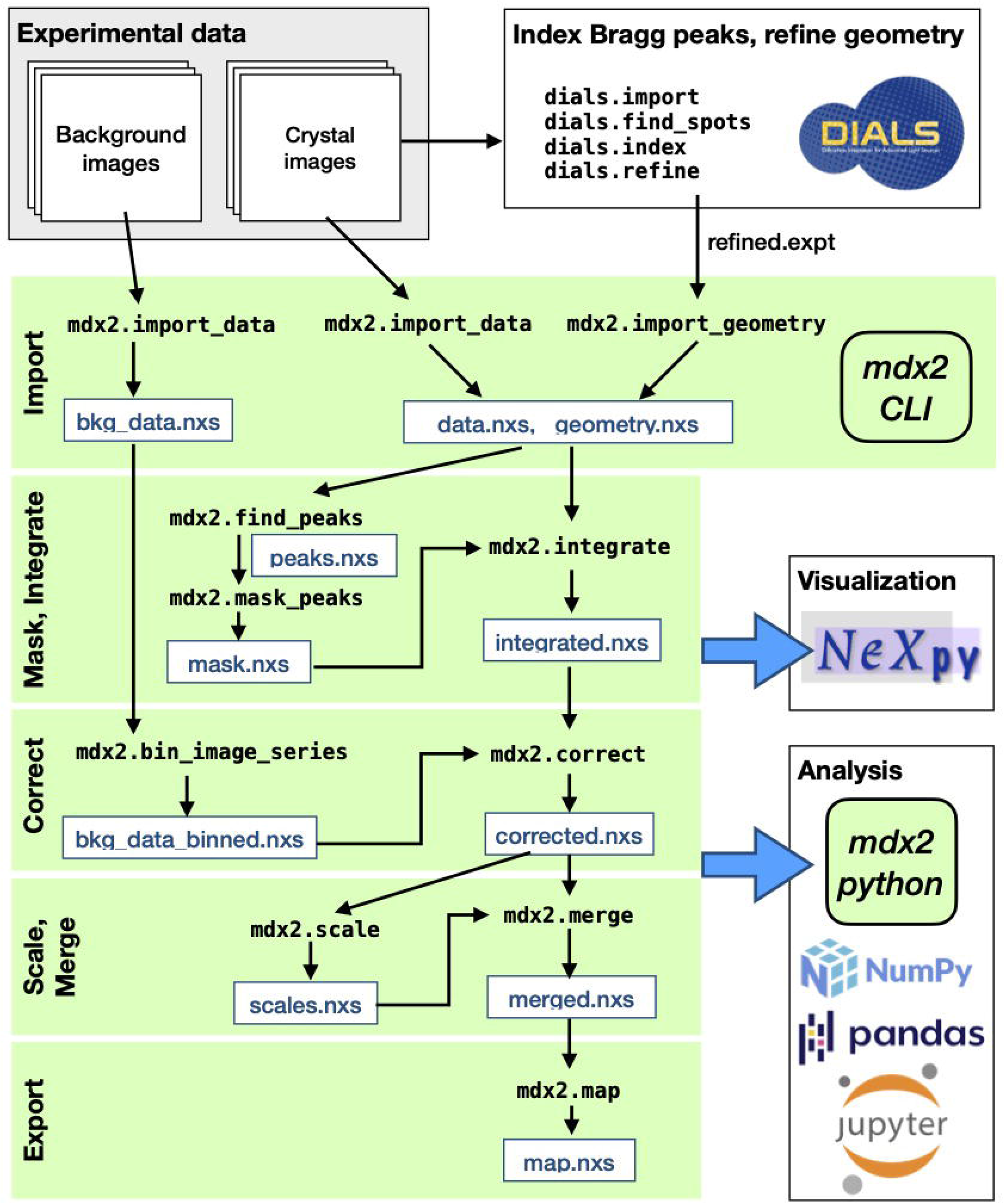
Workflow for reciprocal space reconstruction using *mdx2*. Rotation data from crystals and corresponding background images are collected using procedures described in Chapter 1 (Pei et al., 2023). Bragg peaks in the crystal images are processed using *DIALS* in order to refine the diffraction geometry. Using the *mdx2* command-line interface (CLI), the *DIALS* geometry and diffraction data are imported in NeXus-formatted hdf5 files (.nxs extension) for convenient visualization using *NeXpy*. Subsequent *mdx2* CLI programs (green shading) integrate, correct, scale, and merge the data to produce a three dimensional reciprocal space map (see tutorial in Section 3). At each stage of processing, analysis is performed in Python using *mdx2* together with standard scientific packages (see Section 3.7).

The transition from *mdx-lib* to *mdx2* is not yet complete, and key features have not been implemented at the time of writing. Unlike *mdx-lib, mdx2* does not yet include a full scaling model, and it does not take advantage of multi-CPU architectures. However, *mdx2* is sufficiently capable to make it an excellent tool for data exploration and education, and it is used for the tutorial in Section 3.

## 2. Theory

The goal of reciprocal space mapping is to reconstruct the three-dimensional elastic X-ray scattering signal given many two-dimensional diffraction images of a crystal in different orientations (for example during a ⍰ scan). From a geometric point of view, each pixel in each image must be related to a point in the three dimensional reciprocal space of the crystal. However, geometry is not what makes recirocal space mapping difficult. Instead, the difficulty lies in handling the non-ideal aspects of the data. Compared with the intense Bragg peaks, diffuse scattering is a weak signal because it is spread throughout reciprocal space, and thus in any individual image, the signal may not be visibile above the noise. Diffraction images also contain “background,” either from materials other than the crystal, or from inelastic scattering of atoms within the crystal itself (Compton scattering). Finally, the illuminated crystal volume (and also the background scattering) potentially varies throughout the data collection, and this must be corrected as part of the reconstruction.

Recently, HPC detectors have enabled a data collection stategy of low dose per frame, fine slicing, and high redundancy (see Chapter 1 (Pei et al., 2023)). This strategy is particularly advantageous for reciprocal space mapping. Because the angle increment per frame is small, reciprocal space can be finely sampled without splitting pixels. Furthermore, since many redundant observations are made, the systematic errors can be averaged out or otherwise corrected during scaling and merging.

In Section 2.1, we first review the Ewald construction for visualizing the geometric relationship between detector space and reciprocal space, as this intuition is essential for appreciating the data reduction task. In Section 2.2, we discuss the physics of X-ray scattering and detection needed to relate the number of photons measured to the reciprocal space intensity in electron units. Finally, in Section 2.3, we describe the approach taken by *mdx-lib* to handle the unique features of high-redundancy HPC datasets.

### 2.1 Scattering geometry

In a typical macromolecular crystallography experiment, the crystal is placed in a monochromatic X-ray beam and diffraction images are recorded on a planar detector. First, we consider the geometric relationship between a detector pixel and the reciprocal space of the crystal. The incident X-rays are described by a *wavevector* **s**_0_ in the direction of the beam. Similarly, a scattered wavevector **s**_1_ points from the crystal toward a detector pixel. For the elastic scattering process considered here, the wavevectors have the same length |**s**_0_| = |**s**_1_| = λ^-1^, where λ is the X-ray wavelength. The *scattering vector* **s** is defined as the difference between the incident and scattered wavevectors as follows:

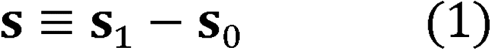

Importantly, the intensity of interest is a function of the scattering vector, *I* = *I*(***s***). Thus, Equation 1 relates the geometry of the experiment (**s**_0_ and **s**_1_) to the reciprocal space of the sample (**s**). This relationship can be visualized intuitively using the Ewald construction (Figure 3A-B).

**Figure 3.**
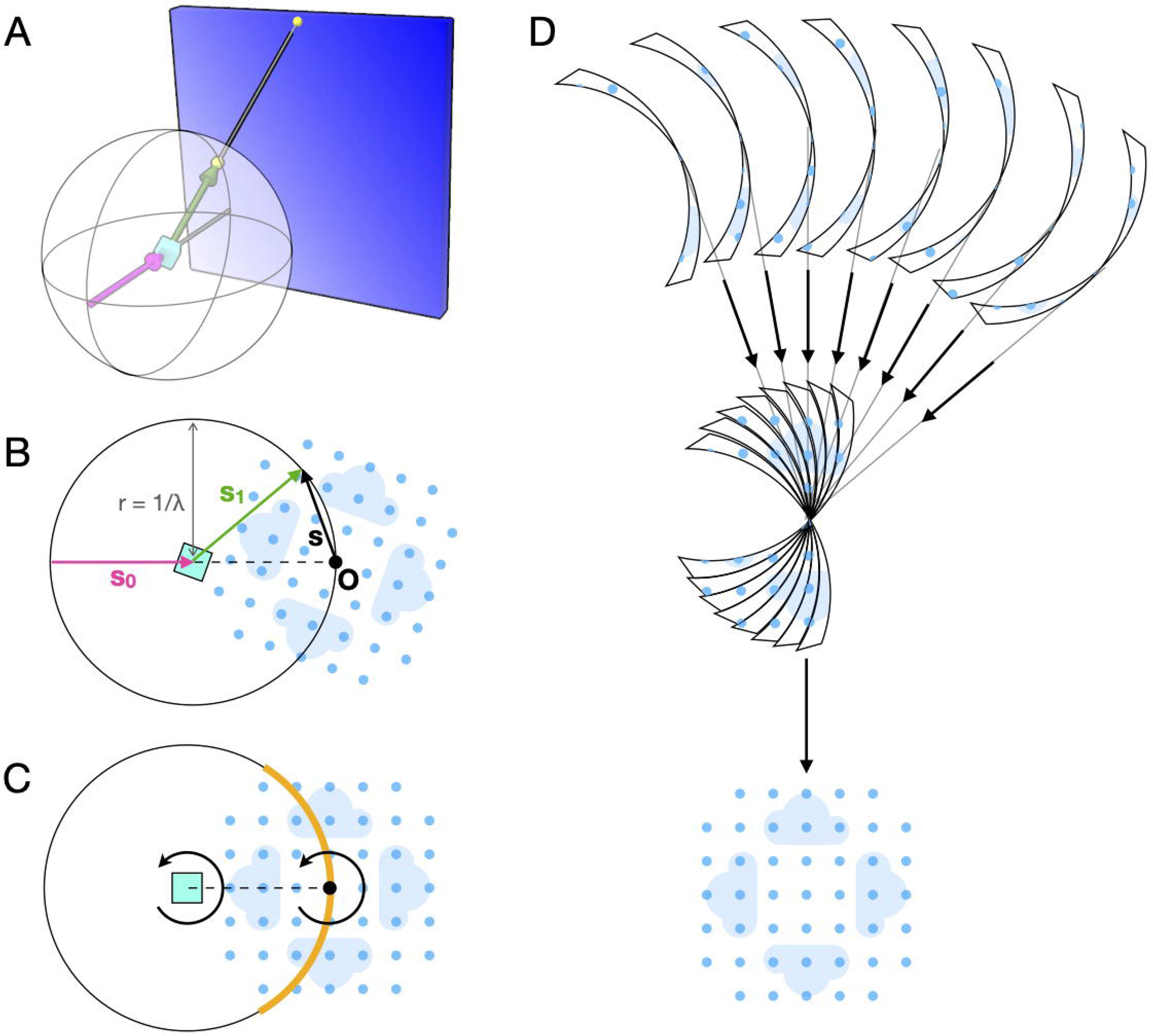
Geometry of reciprocal space reconstruction. (A) An X-ray beam (magenta arrow) is incident on a crystal (cyan) and scattering is recorded on a two-dimensional detector (blue rectangle). Each detector pixel is projected onto the surface of the Ewald sphere, drawn centered on the crystal. The projection (yellow points) is illustrated for a particular scattering direction (green arrow). (B) The same geometry as (A) viewed in the plane formed by the incident and scattered wavevectors (s_0_ and s_1_). In the Ewald construction, the origin of reciprocal space (O) is placed as shown so that the scattering vector s lies on the surface of the sphere. The blue pattern represents a slice through the reciprocal space intensity *I*(s) with lightly shaded clouds representing diffuse scattering and darker circles representing Bragg peaks at the reciprocal lattice points. (C) The pattern of scattering depends on the intersection of *I*(s) with the Ewald sphere (orange region), which rotates as the crystal rotates. (D) Scattering data recorded at different orientations (slices are drawn with finite thickness to illustrate continuous rotation during exposure) are merged in three dimensions to recover the reciprocal space map (first set of arrows), and known symmetries may be used to further complete the reconstruction (bottom arrow).

If the crystal is rotated (for instance during a *ϕ* scan), the reciprocal space coordinate system rotates along with it in the same direction (Figure 3C, arrows). The scattering vector for a particular pixel remains stationary in the laboratory frame, and therefore it samples a different part of reciprocal space. Taken together, the detector pixels map onto a curved surface in reciprocal space, the Ewald sphere (Figure 3C, orange).

Conventionally, the reciprocal space coordinate axes are fixed to the reciprocal unit cell vectors (**a**\*, **b**\*, and **c**\*), such that Bragg peaks have integer coordinates (Miller indices *h, k*, and *l*). Note that for certain space groups the reciprocal lattice vectors are not orthogonal. The change of basis between this *fractional* reciprocal space and the Cartesian, laboratory frame can be written compactly in matrix notation as follows (Busing & Levy, 1967; Waterman et al., 2016):

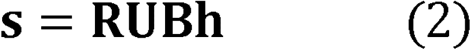

where **h** = (*h,k,l*) is a vector of Miller indices, **B** is a matrix whose rows are the reciprocal unit cell vectors, and **U** and **R** are pure rotation matrices. **U** describes the orientation of the crystal at a particular reference point in the scan (for instance at *ϕ* = 0), and **R** describes the scan-varying rotation (for instance, the rotation about a spindle axis by the amount *ϕ*). The inverse of Equation 2 relates the laboratory coordinates (i.e. a pixel location via Equation 1) to the fractional reciprocal space:

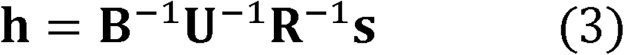

Thus, from a series of images collected with the crystal in different orientations, it is possible to reconstruct the three dimensional reciprocal space (Figure 3D).

Scattering from a crystal obeys certain symmetries according to its space group (Rupp, 2009), and thus the full reciprocal space can often be reconstructed without sampling all possible orientations. Neglecting resonant (anomalous) scattering for simplicity, the symmetry of reciprocal space is described by the crystal’s Laue group, which is the point group of the crystal plus a center of inversion (the point group contains only the rotation operators of the space group; the center of inversion comes about because Fourier transforms of a real function are centro-symmetric). For instance, the space group P 2_1_ has the general positions (*x, y, z*) and (-*x, y* + 1/2, - *z*), with a rotational part corresponding to the point group 2. The corresponding Laue group is 2/*m*, with general positions (*h, k, l*), (-*h*, -*k*, -*l*), (*h*, -*k, l*), and (*h, -k, l*). Thus, only one fourth of reciprocal space is unique (the rest can be generated from the reciprocal space asymmetric unit by applying the Laue group operators). Furthermore, if the crystal is rotated by 360 degrees, each point in reciprocal space is sampled twice (e.g., once by each half of the detector). Neglecting the curvature of the Ewald sphere for simplicity, a 360 degree dataset from a P 2_1_ crystal would be 8-fold redundant (equivalent points are measured 8 times during the scan). Such redundancy can be used to correct for systematic errors because generally the equivalent observations are made by different detector regions and at different crystal orientations (see Section 2.3).

### 2.2. Components of diffraction

As discussed above, a detector pixel records scattered X-rays corresponding to a particular location in the crystal’s reciprocal space. The number of X-ray photons recorded depends on the three-dimensional intensity function of interest, but also on the physics of the scattering process, response of the detector, and background scattering from other sources.

Consider an X-ray image taken as the crystal rotates continuously by a small amount *Δϕ* about an axis 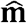 for time *Δt* during which counts are accumulated by a detecor pixel with solid angle ΔΩ. When projected onto the Ewald sphere, the pixel sweeps out a small reciprocal space volume Δ*v*\* during the rotation:

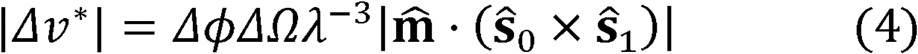

where **ŝ**_0_ and **ŝ**_1_ are unit vectors in the direction of the incident and scattered wavevectors respectively. The intensity recorded by the pixel can be written as an average over the swept volume, defined as follows:

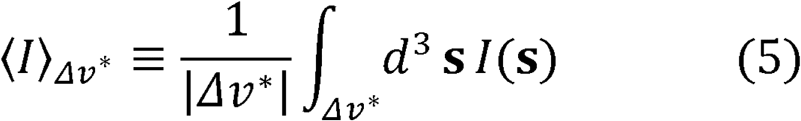

If the intensity varies smoothly (i.e., constant over the exposure) the integral simplifies and the recorded intensity is effectively a sample of *I*(s) at the mean value of **s**, which we call **s**:

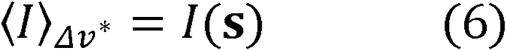

In contrast, consider what happens when a Bragg peak is integrated. The total scattering (Bragg plus diffuse) can be modeled as follows (Meisburger et al., 2017):

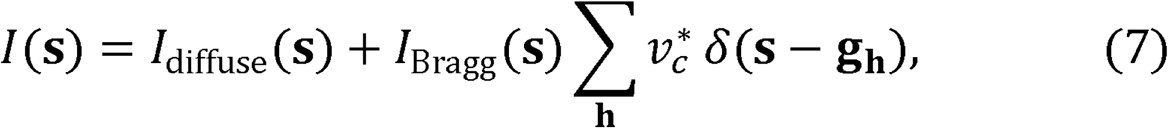

where **g**_h_ are nodes of the reciprocal lattice and 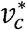 is the reciprocal unit cell volume. Assuming the diffuse scattering varies smoothly and a Bragg peak with Miller index **h** is fully recorded, the average intensity is as follows:

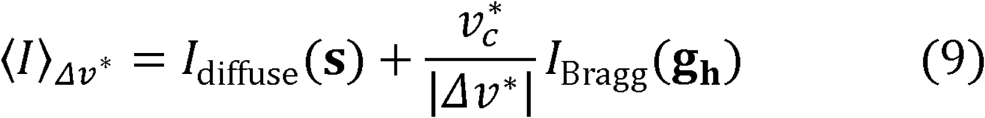

Note that the Bragg intensity is modified by a geometric prefactor (proportional to the Lorentz correction factor). Although Lorentz corrections are applied during Bragg peak integration, they should not be used when correcting diffuse scattering. The Lorentz factor is needed only when features of interest are smaller than the volume of reciprocal space integrated by the detector pixels. In general, it is important to remember that while Bragg and diffuse data processing are similar, there are subtle differences arising from the distinct signal properties. For the purposes of this chapter, we assume Bragg peak integration has already been performed (e.g. using conventional software like *XDS* or *DIALS*), and focus instead on processing the continuous scattering. Note that for Equation 6 to be valid, care must be taken to integrate the reciprocal space map on a fine grid and to exclude pixels containing Bragg peaks from the analysis.

The number of photons recorded by the detector pixel can be calculated based on the geometric considerations, properties of the detector, and the physics of X-ray diffraction. Previous reviews may be consulted for a complete derivation (Meisburger et al., 2020; Meisburger et al., 2017); here we give the result. Let the beam illuminate a volume V_0_ of the crystal with an incident flux (photons per second) *J*_0_. The number of photons recorded *n* is as follows:

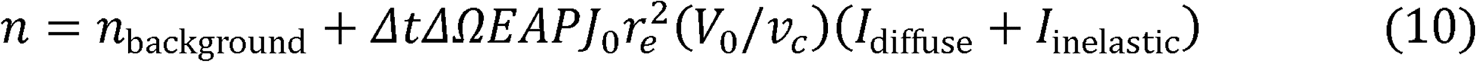

where *n*_background_ are the photon counts due to background, *E* is the efficiency of the detector pixel, *A* is the attenuation factor due to materials between the crystal and the detector, *P* is the polarization factor, *r*_e_ is the Classical electron radius, *v*_c_ is the unit cell volume, and *I*_diffuse_ and *I*_inelastic_ are intensities per unit cell. Thus, to measure *I*_diffuse_, all of the unknown quantities in Equation 10 must be measured or calculated theoretically.

The factors *ΔΩ, E, A*, and *P* can be calculated from the diffraction geometry and known material properties. The solid angle of a pixel is:

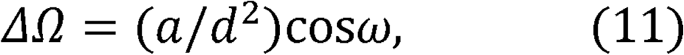

where *a* is the pixel’s area, *d* is its distance from the crystal, and *ω* is the angle between the detector surface normal and the incident X-ray. The pixel’s absorption efficiency depends on the material and thickness *δ* as follows:

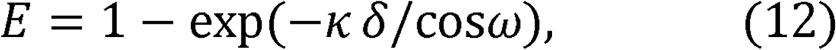

where *κ* is the absorption coefficient (e.g. of Silicon at the X-ray wavelength used). The material between the sample and detector is typically air, whose transmission can be calculated knowing the absorption coefficient *μ* as follows:

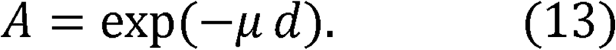

Finally, the polarization effect modifies the scattering efficiency as follows:

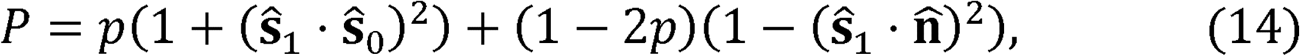

where 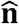 is a unit vector normal to the plane of polarization. Synchrotron X-ray sources are typically polarized in the plane of the ring with a polarization factor *p* ∼ 1.

With geometric factors calculated from first principles, the remaining unknown quantities in Equation 10 are the incident flux, the illuminated volume, the background scattering, and the inelastic scattering from the crystal. These quantities are estimated in later stages of data processing, for instance by utilizing the redundant observations to derive relative scale factors for different diffraction images (see Section 2.3). In the mean time, an intermediate quantity *I*_measured_ is computed from the corrected photon counts as follows:

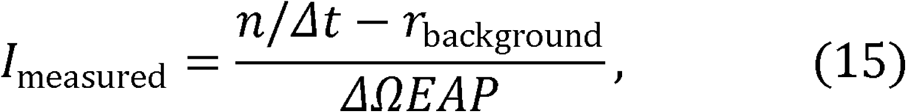

where *r*_background_ is the background scattering rate, which has been measured separately, e.g. by translating the crystal out of the beam (see Chapter 1 (Pei et al., 2023)).

### 2.3. Estimating and correcting errors

Error estimation is an essential part of data processing, and it begins during integration. After masking out the Bragg peaks, pixels from adjacent images are grouped together according to which voxels they contribute to in the (unsymmetrized) map, and their counts are summed. The uncertainty due to Poisson statistics for an observation with *n* photon counts is √*n*. This uncertainty is propagated through all subsequent correction steps (Equation 15) so that each measured intensity value has a corresponding error estimate. The errors are used as weighting factors for scaling, as described below.

Typically, each voxel (or its symmetry equivalent) is observed multiple times in the dataset at different points in the scan. This redundancy can be used to estimate unknown scale factors in Equation 10. The scaling algorithm minimizes the least-squares difference between *I*_predicted_ and *I*_measured_ weighted by the uncertainty estimates (σ):

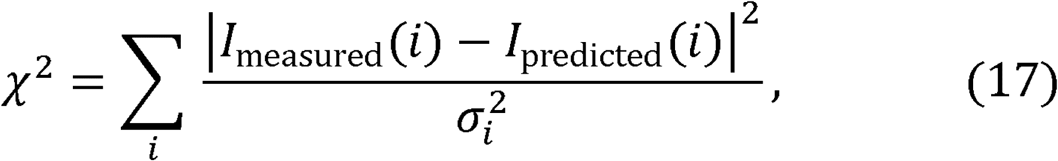

where the sum runs over all observations *i. I*_predicted_ comes from the scaling model: it is a function of the scaling parameters and of the merged intensities *I*_merged_. Initially, both the scale factors and the merged intensities are unknown, and both must be refined simultaneously. In general, the optimization problem is nonlinear but it can be solved by an alternating least squares (ALS) algorithm (Hamilton et al., 1965), which alternates between two steps: merging the data with scale factors fixed, and determining scale factors given the merged data.

Let the scaled (but not merged) intensities and uncertainties be *I*_scaled_ and σ_scaled_. Assuming a linear relationship between *I*_measured_ and *I*_scaled_ (which is a general feature of scaling models for X-ray diffraction), Equation 17 can be rewritten as follows:

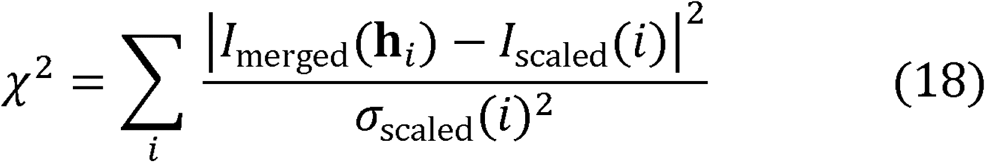

where **h**_*i*_ refers to the index of the observation mapped to the asymmetric unit of reciprocal space. Let the index *j* refer to all observations mapping to a particular voxel **h**. The value of *I*_merged_ that minimizes Equation 18 is as follows:

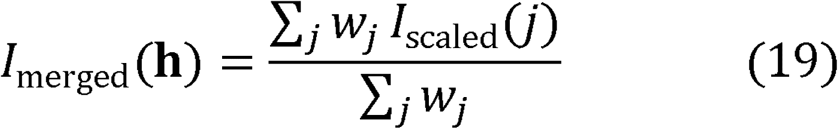

where 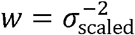 The corresponding uncertainty in the merged intensity is estimated as follows:

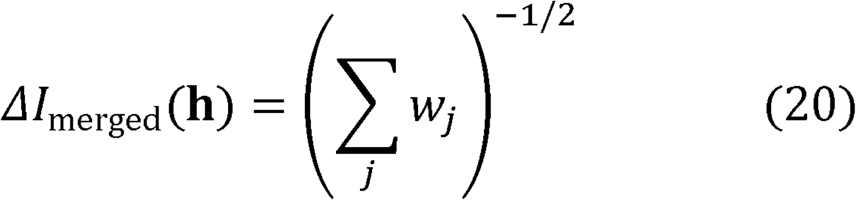

In *mdx-lib*, the following scaling model is refined:

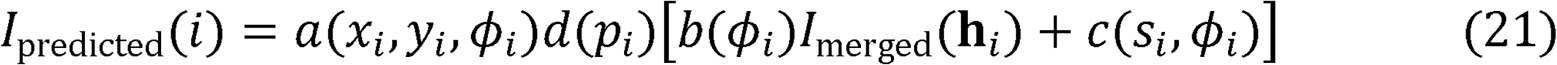

The model relates the merged intensities for a certain voxel in the scaled and merged map *I*_merged_(**h**_*i*_) to a predicted value in the unscaled, unmerged dataset *I*_predicted_(*i*). Each observation has three scale factors (*a, d*, and *b*) and one offset term *c*. The scale factor *a* accounts for self-absorption of the crystal, and its value depends on the position of the observation on the detector (*x*_i_, *y*_i_) and the rotation angle of the crystal *ϕ*_i_. The scale factor *d* corrects for detector flat field errors, and its value depends on which detector chip recorded the counts (index *p*_i_). The scale factor *b* depends only on the rotation angle, and it corrects for changes in illuminated volume and incident beam intensity during the scan.

Programs for scaling Bragg data, such as Aimless (Evans & Murshudov, 2013) and XDS (Kabsch, 2010), include scale factors similar to those in Equation 21. However, the offset term *c* is unique to diffuse scattering. Its purpose is to compensate for extra background scattering that is not included in the background measurement, for instance from liquid on the crystal surface or the loop materials. The scaling model assumes that the added background is isotropic (varying as a function of resolution, *s*_i_). When the scaling model is refined, the value of *c* is restrained so that it is always positive (or zero).

The scaling algorithm solves for the unknowns in Equation 21 by minimizing ***χ***^2^ (Equation 17), while also enforcing smoothness of the scale factors to avoid overfitting (Meisburger et al., 2020). The terms in the scaling model are physically-motivated, and thus after refinement the scale factors should be inspected to verify that they make physical sense (Figure 4).

**Figure 4.**
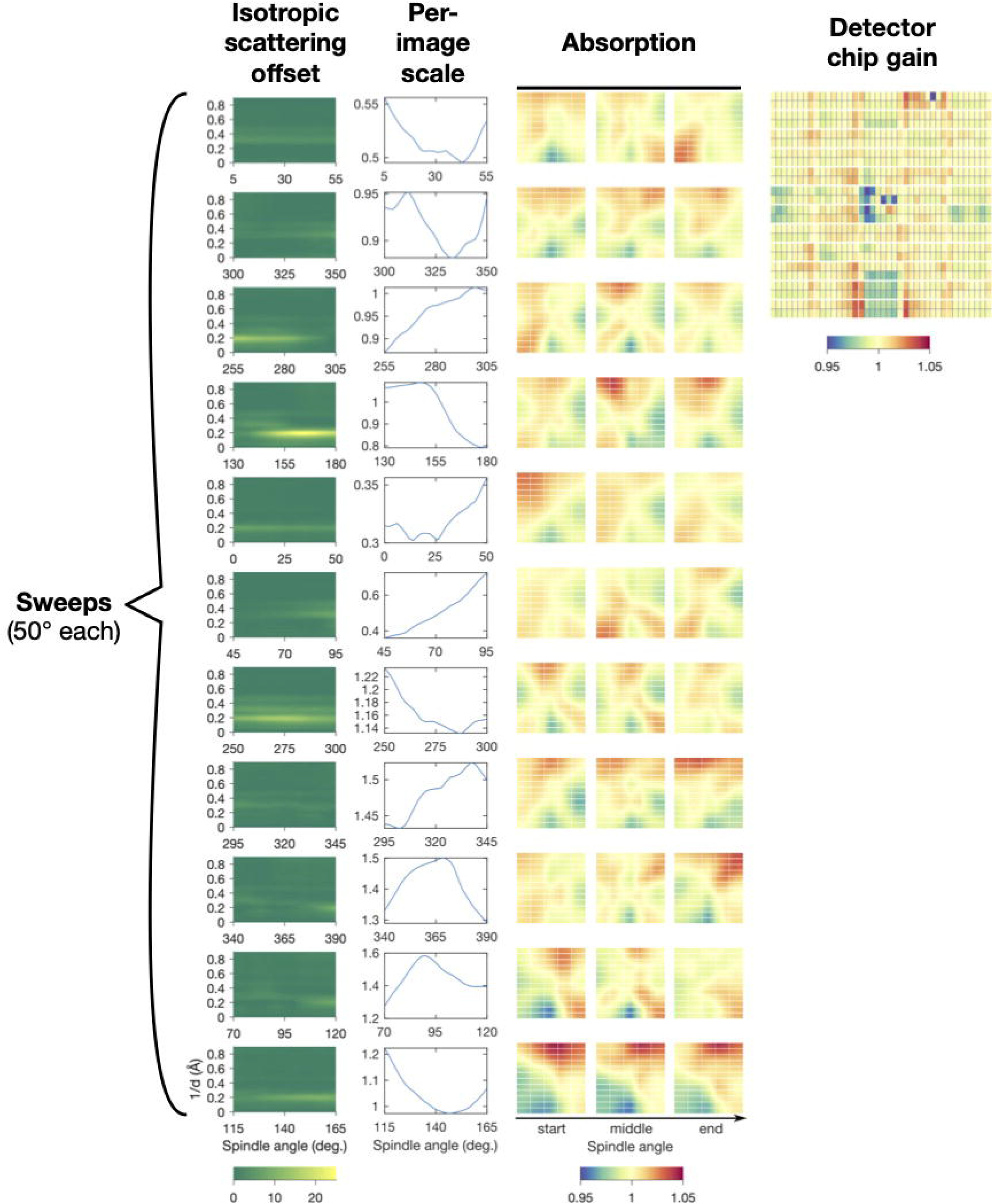
Scaling model for a triclinic lysozyme dataset from refinement using *mdx-lib*. Panels are arranged with rows corresponding to the eleven sweeps of data distributed over four crystals (see original publication for details) and columns corresponding to the four terms in the scaling model in Equation 21 (left to right): *c, b, a*, and *d*. Reproduced under CC-BY-4.0 license from (Meisburger et al., 2020).

By refining the scaling model, the intensities of redundant observations are brought into agreement, however the absolute scale factor has not been determined (i.e., the product of photon flux and illuminated volume in Equation 10). Depending on the application, it may be necessary to place the map on an absolute scale; for instance, it is required to subtract a simulated inelastic term in Equation 10. We previously developed a method based on comparing the simulated and experimental total scattering (Meisburger et al., 2020, 2023) which has been implemented in *mdx-lib* (for examples, see https://github.com/ando-lab/mdx-examples).

## 3. Data Processing Tutorial

To provide a concrete example of the steps involved in data processing outlined above, we present an introductory tutorial using the *mdx2* software. The tutorial can be completed by following instructions and typing commands in this chapter, or by downloading and stepping through the accompanying Jupyter notebooks (see Appendix A.1). A relatively small dataset was chosen for this tutorial so that it can be completed on a personal computer (specific requirements are given in Appendix A.1). A room-temperature X-ray diffraction dataset from bovine insulin in space group I2_1_3 (Figure 5) was collected at the Cornell High Energy Synchrotron Source (CHESS) beamline F1 following protocols described previously (Meisburger et al., 2020, 2023) and summarized in Chapter 1 (Pei et al., 2023). The insulin dataset consists of a ⍰ scan with 500 diffraction images (0.1 degrees each, 50 degrees total) and 50 background measurements (1 degree each over the same angular range). The dataset is available online (see Appendix A.2).

**Figure 5.**
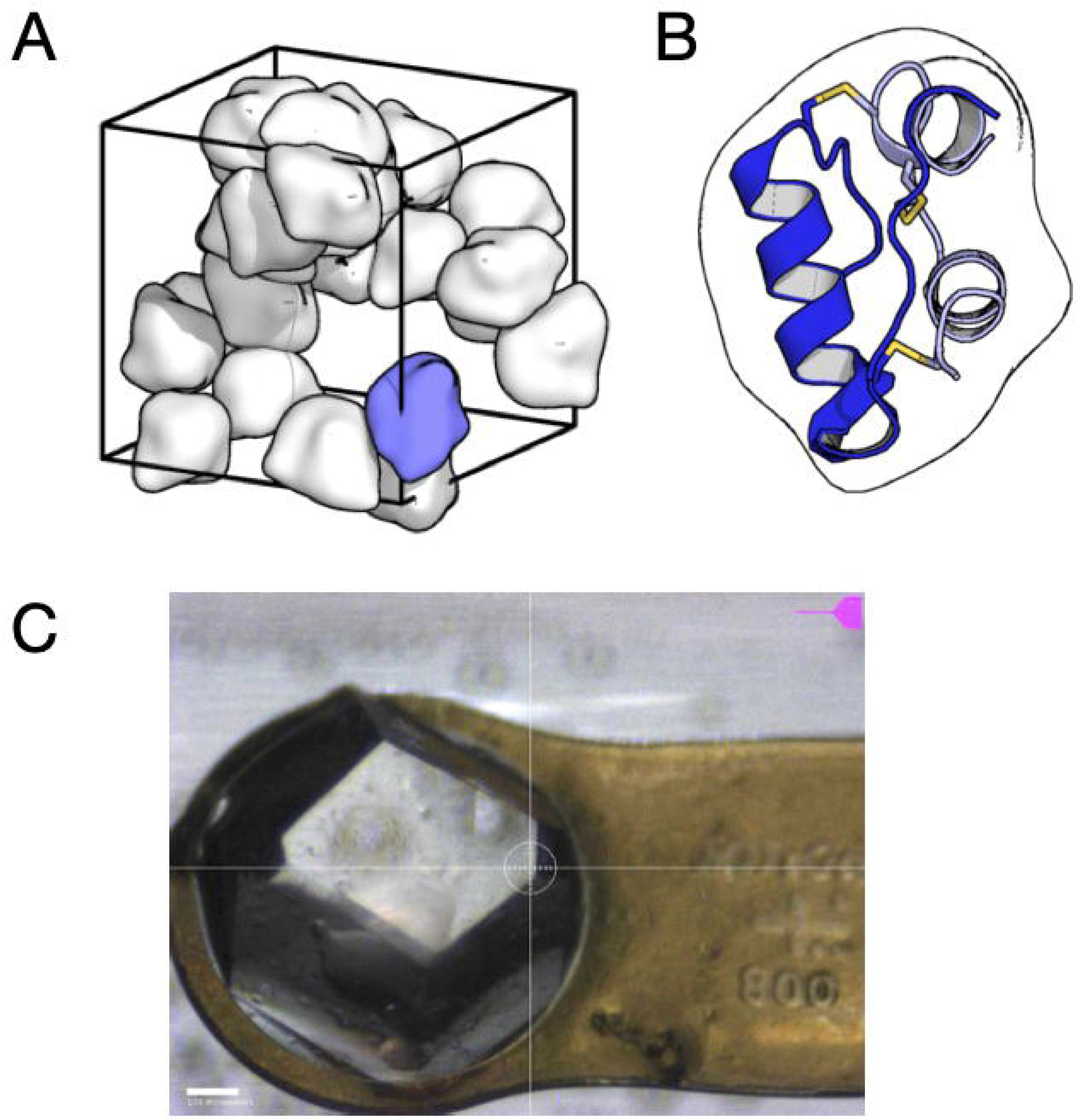
Insulin crystal that produced the dataset used in the tutorial. (A) Insulin from *Bos taurus* was crystallized using a Zn-free condition in the space group I2_1_3. The unit cell (wireframe box) contains 24 copies of the asymmetric unit (smooth molecular surfaces). The packing arrangement has large channels occupied by disordered solvent. (B) The asymmetric unit, drawn in the same orientation as the blue blob in (A), contains a single insulin molecule consisting of two alpha-helical chains (blue and gray cartoons) cross-linked by three disulfide bonds (yellow sticks). (C) Image taken during data collection of the ∼800 micron crystal mounted on a Kapton loop and held within a plastic capillary to maintain hydration. The datset processed in this tutorial consists of a single 50-degree sweep (0.1 degrees and 0.1 seconds per frame). Because of the large crystal size and high symmetry, a single sweep is sufficient for a complete reciprocal space map.

The first step is to install the software. The recommended method is to create a stand-alone Python environment using *miniconda3* (Appendix A.3) and then install *mdx2* from the GitHub repository using *pip* (Appendix A.4). In addition to *mdx2*, this data processing tutorial uses *DIALS* for pre-processing, *NeXpy* for visualization, and *Jupyter* for interactive notebooks. These programs can be installed and run separately, but for convenience we recommend installing them in the same Python environment as *mdx2* (Appendix A.5).

### 3.1 Indexing and Geometry Refinement

The first step in data processing is to determine the diffraction geometry and crystal orientation by analyzing the locations of strong Bragg peaks. These steps are performed using *DIALS*, a suite of programs for integrating and scaling Bragg data (Winter et al. (2022)). For our purposes, we will only use indexing and geometry refinement tools. Normally these steps would be followed by integration and scaling, but we will skip them to save time. Detailed tutorials are available on the *DIALS* website (https://dials.github.io).

#### Metadata import

The image file headers contain metadata specifying the experimental geometry such as X-ray wavelength, detector distance, oscillation increment per image. The command `dials.import` reads the image headers and outputs a json-formatted text file `imported.expt` containing the relevant metadata. In the terminal, change to the working directory (containing the `images` folder) and run the following command to import the insulin dataset:

~~~
dials.import images/insulin_2_1
~~~

#### Beamstop mask

The beamstop mask must also be generated and added to the `imported.expt`. A beamstop mask is not essential for Bragg data processing, but it is important to create one for later import into *mdx2*. The mask can be drawn using the graphical image viewer that comes with *DIALS*:

~~~
dials.image_viewer imported.expt
~~~

Alternatively, the following will generate an appropriate circular mask for the insulin dataset using `dials.generate_mask` and `dials.apply_mask`.

~~~
dials.generate_mask imported.expt untrusted.circle=1264,1242,50
dials.apply_mask imported.expt input.mask=pixels.mask
output.experiments=imported.expt
~~~

Note that the beamstop shadow mask is relatively simple in this case because the beamstop was suspended on an X-ray transparent Kapton film. It is more common to mount the beamstop on a rigid support whose shadow must be more carefully masked.

#### Spotfinding

Next, `dials.find_spots` will read all of the images and locate Bragg peaks. This requires reading in all of the diffraction images, and it can take a while depending on computational resources.

~~~
dials.find_spots imported.expt
~~~

#### Indexing

The indexing step assigns Miller indices to each diffraction spot. When processing an unknown crystal, it is usually necessary to re-index the data after space group determination. Here, the space group is known (the insulin crystal is I2_1_3, space group number 199), and we can avoid having to re-index later by passing this information to `dials.index`.

~~~
dials.index imported.expt strong.refl space_group=199
~~~

For diffuse scattering, it’s important to verify that all (or nearly all) peaks are indexed. There might be split Bragg reflections, multiple lattices, or salt crystals contaminating the signal. Or, the crystal might have slipped during data collection.

Check that indexing was successful by examining the *dials*.*index* output.

Find the table with “% indexed.”

| Imageset | # indexed | # unindexed | % indexed |
| --- | --- | --- | --- |
| 0 | 50694 | 417 | 99.2% |

The fraction of indexed peaks should be close to 100%. A small percentage may suggest incorrect experimental geometry, multiple lattices, twinning, salt or ice diffraction, or other issues. The indexing results should be inspected by running `dials.image_viewer indexed.expt`. In our case, 99.2 percent of strong peaks were indexed (there were 417 unindexed peaks and 50694 indexed peaks).

Find the table called “RMSDs by experiment.”

`RMSDs by experiment:`

| Exp id | Nref | RMSD_X (px) | RMSD_Y (px) | RMSD_Z (images) |
| --- | --- | --- | --- | --- |
| 0 | 4999 | 0.21024 | 0.2213 | 0.20059 |

This table includes the root mean square displacement (RMSD) between the predicted and observed spot centroids in units of pixels (in *x, y* directions) and images (in *z* direction). If the RMSDs are less than ∼1, this indicates that the indexing solution was successful and a single set of geometric parameters can describe the entire dataset accurately. However, if the crystal slips slightly during data collection or the lattice constants change, the spot predictions can be improved by fitting a scan-varying geometric model.

#### Geometry refinement

Next we use `dials.refine` to fit a scan-varying model of the crystal geometry. This can be important if the lattice constants change due to radiation damage, or if the crystal is not held rigidly in the loop and slips during data collection.

~~~
dials.refine indexed.expt indexed.refl
~~~

To see if the spot predictions improved, find the table “RMSDs by experiment” in the text output:

`RMSDs by experiment:`

| Exp<br>id | Nref | RMSD_X<br>(px) | RMSD_Y<br>(px) | RMSD_Z<br>(images) |
| --- | --- | --- | --- | --- |
| 0 | 45511 | 0.17561 | 0.18017 | 0.19491 |

Compared with the corresponding table from `dials.index`, the RMSDs have improved as we expected. However the improvement is modest. In our case, the indexing solution already had small RMSDs and refinement was not strictly necessary.

#### Diagnostic statistics and plots

The `dials.report` function generates an html file with interactive plots. The report should be examined closely to catch any issues that would potentially complicate analysis of the diffuse scattering.

~~~
dials.report refined.expt refined.refl
~~~

Open the file `dials.report.html` in an internet browser.

Expand the tab called “Analysis of scan varying model” (Figure 6A,B). These plots should be checked to see whether the unit cell axes change significantly during data collection. Abrupt changes in the orientation parameters may indicate that the crystal slipped during data collection.

**Figure 6.**
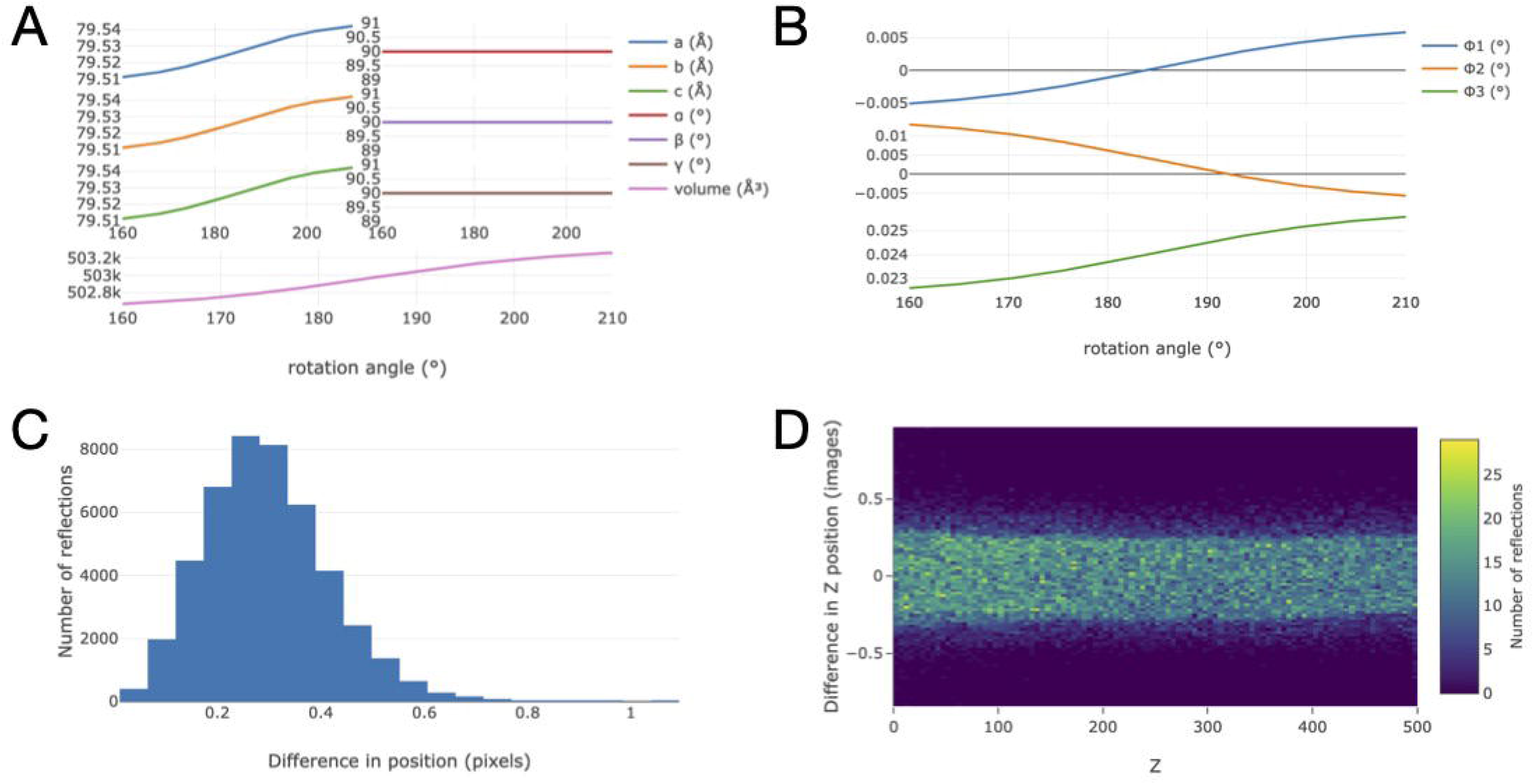
Plots from the *DIALS* report that illustrate a successful geometry refinement. The scan-varying model allows for the unit cell constants (A) and setting angles (B) to vary smoothly throughout the scan. Refinement minimizes the difference between observed and predicted coordinates of the Bragg peaks on the detector surface (C) and image number *Z* in the scan (D). The parameter changes are minimal during the scan, and the errors are much less than one pixel or image. Furthermore, there are no sudden jumps or outliers in panel (D), which might indicate that the crystal had slipped or cracked during data collection.

Expand the tab called “Analysis of reflection centroids.” Ideally, the error in *x* and *y* positions should appear random across the detector face. In our dataset, the errors cluster in rectangular blocks corresponding to the detector modules. If sub-pixel accuracy is desired, the module alignments can be calibrated. Next, examine the plots of spot centroid errors (Figure 6C,D). The spread of centroids around the mean should show a compact peak. If there are multiple peaks observed, then the crystal may be twinned or multiple lattices may be present (e.g. due to cracking).

#### Background image metadata

An additional dataset was acquired for background subtraction in *mdx2*. We’ll use `dials.import` and `dials.apply_mask` to create a metadata file `background.expt` that will be used in Section 3.4.

~~~
dials.import images/insulin_2_bkg output.experiments=background.expt
dials.apply_mask background.expt input.mask=pixels.mask
output.experiments=background.expt
~~~

### 3.2 Data Exploration

In this section, we prepare data files for reciprocal space mapping with *mdx2* and demonstrate interactive tools for data exploration.

#### Importing diffraction data

First, we introduce the *mdx2* command line programs for importing diffraction images and geometric corrections from *DIALS* in Section 3.1. Most features of *mdx2* needed for the tutorial are available in command-line programs. Run the following program to verify that *mdx2* is installed and activated:

~~~
mdx2.version
~~~

The version number (0.3.1) should be printed.

The first step in the *mdx2* workflow (Figure 2) is to import the diffraction images. Unlike *DIALS, mdx2* does not read diffraction data directly from image files. Instead, the data are imported (i.e. copied) into a three-dimensional array (image stack) and stored in a NeXus-formatted hdf5 file. There are two advantages to this: first, subsequent *mdx2* programs can be very simple because they don’t need to handle diverse image formats, and second, the data can be compressed in a way that improves performance when it is subsequently read.

From within the directory containing the *DIALS* output files (`refined.expt`), run the following:

~~~
mdx2.import_data refined.expt --chunks 20 211 493
~~~

The program reads image files (the file path is stored in `refined.expt`) and saves the raw data and associated metadata in the file `data.nxs`. The `--chunks` parameter controls the size of the sub-arrays that will be compressed in the hdf5 file. Chunking allows for reading or writing small segments of the array without decompressing the entire dataset. Here, we chose a size of 20 × 211 × 493. Roughly, 211 × 493 corresponds to a panel on the Pilatus 6M, and 20 specifies a stack of 20 frames (2 degrees of rotation). The particular chunk shape is not critical, but it can affect performance of subsequent operations.

Note that operations involving raw image data can be time consuming. Data import may take several minutes. For slow operations, *mdx2* produces print statements to track its progress. In this case, it prints the frame number as files are being read (0 – 499). Every 20 frames, the program writes chunks incrementally to the data file (`data.nxs` will increase in size until it reaches approximately 1.1 Gb).

The contents of any nexus file can be viewed using the program `mdx2.tree`. For instance,

~~~
mdx2.tree data.nxs
~~~

prints the contents of the data file (Figure 7A).

**Figure 7.**
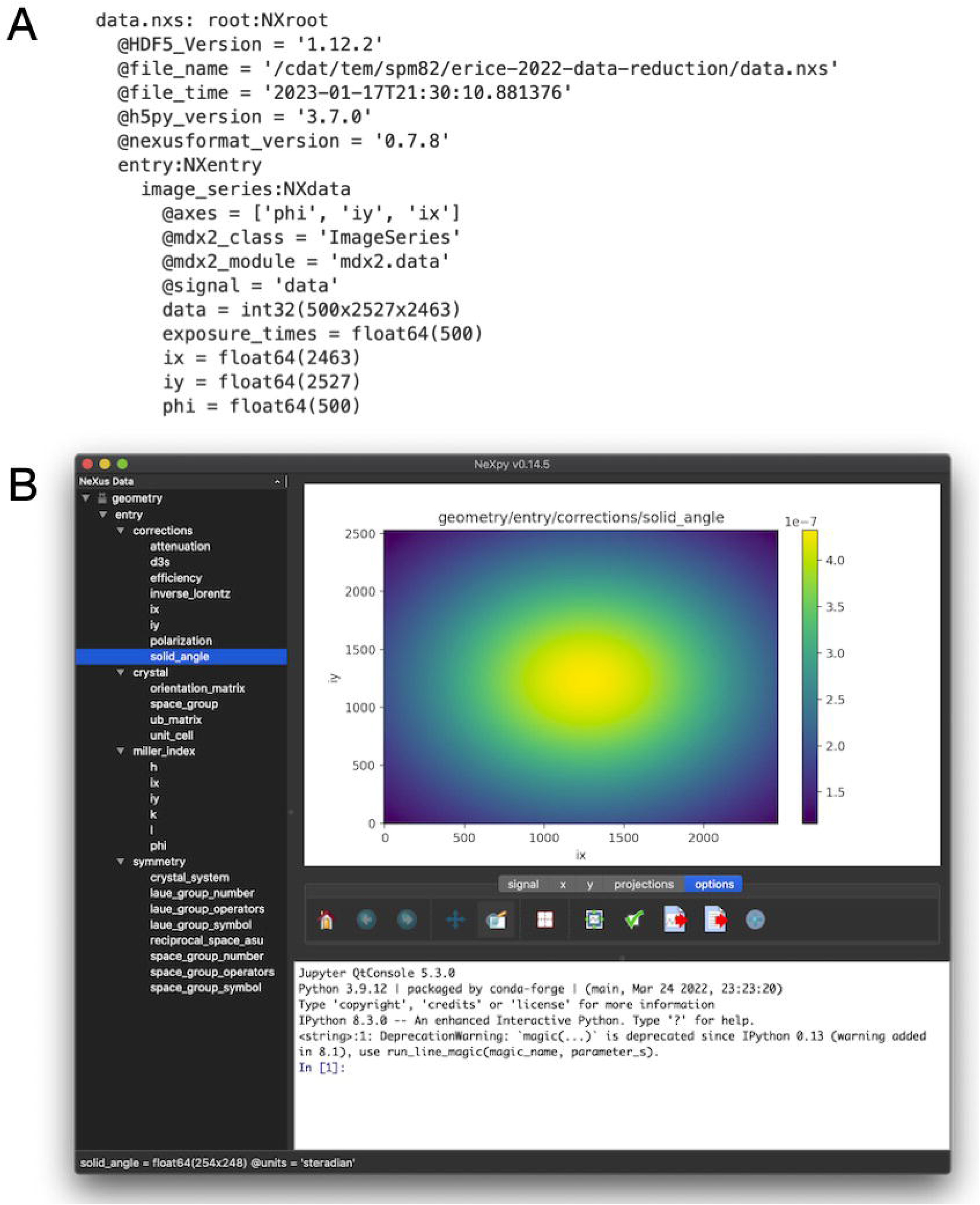
NeXus files produced by *mdx2* and visualization using *NeXpy*. (A) The file `data.nxs` contains the raw diffraction data as a 3-dimensional array with associated metadata in an NXdata object. The text representation of the file was produced by the function `mdx2.tree`. (B) The more complex file `geometry.nxs` has many NeXus objects organized in a hierarchical fashion (tree view, left panel). Objects can be plotted by double-clicking (here, the `solid_angle` correction mapped over the detector surface). Axes are generated automatically from the metadata.

#### Importing diffraction geometry

The next step is to import the geometric information necessary to map each pixel to its location in reciprocal space. Run the following program:

~~~
mdx2.import_geometry refined.expt
~~~

The output file `geometry.nxs` contains crystal lattice information, symmetry operators, and pre-computed arrays of correction factors and Miller indices. To conserve disk space, the array values are not computed at every pixel, but instead on a coarser grid. By default, the corrections are sampled every 1 degree of rotation and every 10 pixels in the *x* and *y* directions. If needed, these parameters can be changed using the optional `--sample_spacing` argument. Help text for any `mdx2` program can be printed by adding `--help`. For instance, running `mdx2.import_geometry -- help` will print the following:

~~~
usage: mdx2.import_geometry [-h] [--sample_spacing PHI IY IX]
                            [--outfile OUTFILE]
                            expt
Import experimental geometry using the dxtbx machinery
positional arguments:
  expt                   dials experiments file, such as refined.expt
optional arguments:
  -h, --help             show this help message and exit
--sample_spacing PHI IY IX
                         inverval between samples in degrees or pixels
                         (default: [1, 10, 10])
--outfile OUTFILE        name of the output NeXus file (default: geometry.nxs)
~~~

Under “optional arguments” above, the help text explains that the default interval for sampling is every 1 degree of rotation and every 10 pixels on the detector.

The contents of `geometry.nxs` can be viewed using `mdx2.tree` as above. However, the output is limited to the metadata such as the array types and sizes. Much more information is available using *NeXpy*. For instance, to view the solid angle correction, follow these steps:

- In the terminal type `nexpy` to launch the program
- Click `File > open`, navigate to your working directory
- Open `geometry.nxs`
- Expand the tree in the left panel to locate `solid_angle`.
- Double-click it to plot.

The solid angle per pixel (steradian units) should be displayed as a colormap on the detector surface (Figure 7B).

It can be educational to browse the data objects in `geometry.nxs` as many of the quantities introduced in Section 2 are pre-computed during the import step. Crystal geometry objects include the `orientation_matrix` (U in Equation 2), `ub_matrix` (product UB in Equation 2), and Miller index maps *h, k*, and *l* (computed using Equation 3 from the *DIALS* scan-varying model). The 3×3 matrices in `laue_group_operators` represent general positions of the Laue group, the 3×4 matrices in `space_group_operators` represent general positions of the space group operators (the extra column is needed to represent translations (Rowicka et al., 2004)), and `reciprocal_space_asu` is a mathematical expression defining the asymmetric unit of reciprocal space. Correction factors introduced in Section 2.2 include `solid_angle` (Equation 11), `efficiency` (Equation 12), `attenuation` (Equation 13), `polarization` (Equation 14), `d3s` (Equation 4), and `inverse_lorentz` (the absolute value factor in Equation 4).

#### Calculating a rocking curve in Python

Finally, we demonstrate how saving diffraction data in NeXus files can enable rapid data exploration in Python. The following lines in Python are all that is needed to plot a rocking curve for an intense reflection (Figure 8A):

**Figure 8.**
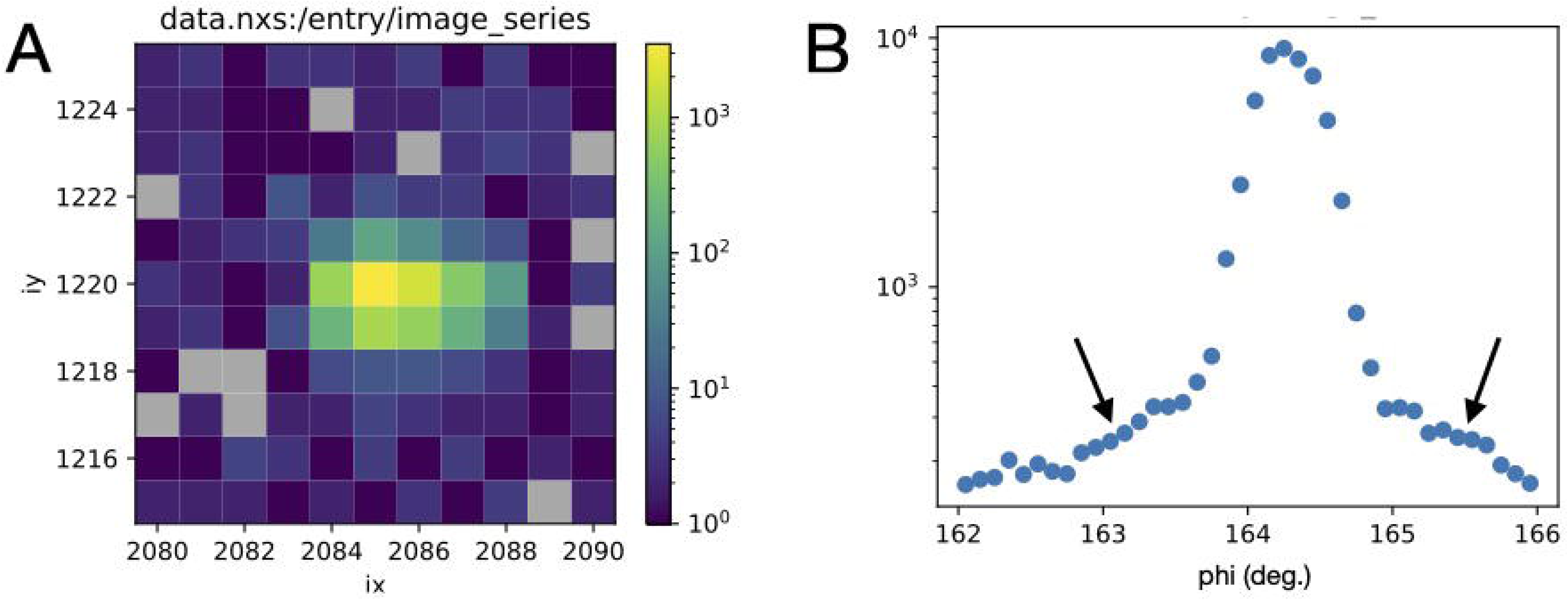
Rocking curve showing Bragg diffraction and diffuse scattering. An intense peak was selected that passed slowly through the Ewald sphere (lying close the rotation axis in reciprocal space). (A) The Bragg peak in frame 42 where it’s intensity is maximum. The intensity decays quickly to background over several pixels (note the logarithmic color scale; pixels with zero counts are colored gray). (B) The total number of counts within the region around the Bragg peak in (A) as a function of rotation angle. The central bell-shaped peak is the Bragg diffraction, and the gradual tails are diffuse halos (arrows). The plots were generated using built-in functions of the *nexusformat* package (see Section 3.2).

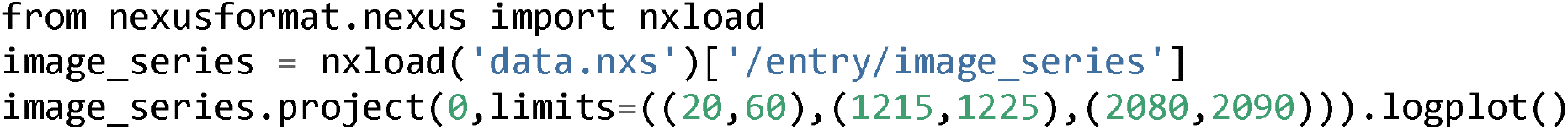

Here, *project* is a method associated with the NXData object called `image_series` that calculates a projection of a high dimensional array to a subset of the axes. Here, the pixels surrounding the Bragg peak are summing onto axis 0 of the dataset (frame number). The range of pixels considered is given by the `limits` argument: frames 20 through 60, pixels 1215 to 1225 in the *y* direction, and 2080 to 2090 in the *x* direction. A plot is produced similar to Figure 8B. Further examples of working with nexus files in Python can be found in the *nexusformat* documentation (http://nexpy.github.io/nexpy/pythonshell.html).

*Mdx2* does not currently provide a means of locating particular Bragg peaks of interest based on their Miller indices. To find a particular Bragg spot, we recommend overlaying the spot predictions with Miller index labels on diffraction images using *dials*.*image_viewer*.

### 3.3. Bragg Peak Masking

Because different corrections are needed when integrating Bragg peaks and diffuse data (see Equation 9), it is important to ensure that the signals are separated before processing. It is important to note that a Bragg peak’s extent in *ϕ* can vary depending on the distance from the spindle axis in reciprocal space. And, Bragg peaks may not be perfectly symmetric Gaussian peaks. Peak width depends on crystal mosaicity, and this can even vary during a scan. In addition, the diffraction geometry may not be modeled exactly right, leading to further errors in peak position. A masking function must account for all of these effects.

Designing a perfect Bragg peak mask is a tricky problem, and different researchers have approached it various ways. With *mdx2* we take a simple empirical approach. Here is the algorithm:

- List all of the pixels in the image stack above a certain count threshold (“hot pixels”).
- Index all of the hot pixels, converting their position in image space to Miller index (*h, k, l*).
- Subtract the nearest integer from each Miller index leaving only the fractional part (*Δh, Δk, Δl*) which varies from -0.5 to 0.5. Because peaks have finite extent, the values of *Δh, Δk, Δl* will not all be zero, but will look like a cloud of points (Figure 9A).
- Fit all of the points to a single anisotropic 3D Gaussian distribution (Figure 9B).
- Create a binary mask for all pixels in the image stack that excludes pixels falling within the 3D Gaussian region according to a sigma threshold (Figure 9C), and any hot pixels falling outside these regions.

**Figure 9.**
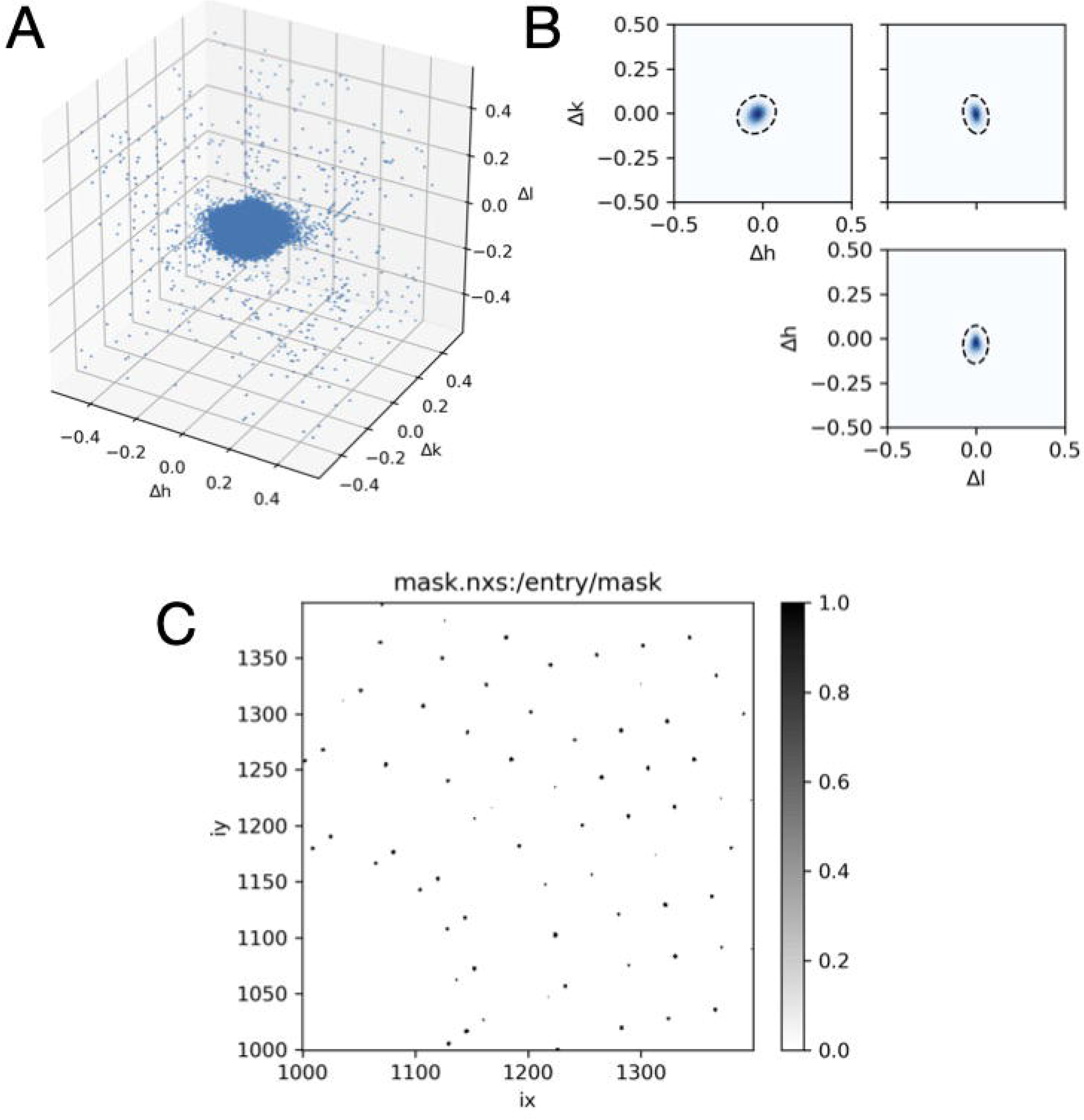
Bragg peaking masking. (A) Strong pixels (those exceeding a count threshold) are mapped into reciprocal space and plotted as a function of distance from the nearest Bragg peak (axes are fractional Miller indices). (B) The points are fit to a three-dimensional Gaussian probability distribution, here shown as a dashed contour line at 3 standard deviations superimposed on orthogonal projections of the distribution in (A). The Gaussian fit accouts for potential anisotropy due to crystal properties and errors in the model of diffraction geometry. (C) A mask is generated for all pixels in each image according to the Gaussian fit.

To execute the masking altorithm in *mdx2*, first perform the strong pixel search:

~~~
mdx2.find_peaks geometry.nxs data.nxs --count_threshold 20
~~~

A threshold of 20 was chosen to be 10 times the background level of ∼ 2 photons / pixel, which works well in this case. *Mdx2* does not currently have a method to automatically estimate the threshold. The images should be inspected manually to determine an appropriate value. When the search finishes, a Gaussian is automatically fit to the point cloud:

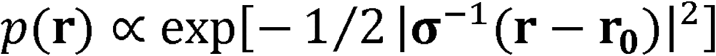

where .**r**=(Δ*h*, Δ*κ*, Δ*l* The best-fit values according to mdx2.find_peaks printed and saved to a file (peaks.nxs):

~~~
6602 peaks were rejected as outliers
GaussianPeak model: r0 = [-0.033588 -0.006928 -0.002245]
sigma =
[[-0.025445 -0.025525 -0.001434]
 [-0.029339 0.021216 0.004702]
 [0.004484 -0.006027 0.022625]]
~~~

Note that the peak center **r**_0_ is not exactly zero, which may indicate a slight error in the diffraction geometry. This error, as well as the extent of the point cloud (*σ*), should be taken into account when designing a reciprocal space grid for integration. The sampling should not be finer than the geometric errors allow. Here, the Bragg peaks are well localized; the width of the Gaussian distribution is approximately 1/20 of the reciprocal unit cell dimension (∼two times the diagonal elements of *σ*, above). Thus, a sampling rate perhaps as fine as 20 × 20 × 20 voxels per Bragg peak could be justified. Note that such fine maps consume a lot of computer memory, and thus for the purposes of this tutorial, a coarser map will be used. In practice, there is not a one-size-fits-all method to determine the optimum sampling as it depends on the signal-to-noise needed and the application (e.g., fitting a power law requires that there should be at least 3 points – ideally more – between the Bragg peak and the edge of the Brillouin zone).

Next, create the mask:

~~~
mdx2.mask_peaks geometry.nxs data.nxs peaks.nxs --sigma_cutoff 3
~~~

The `sigma_cutoff` parameter controls the size of the masked region around the Bragg peak. A value of 3 means that pixels are masked within 3 standard deviations of the peak center according to the model fit above. The function also masks outliers flagged previously. The resulting masks are saved as an image stack in `mask.nxs` and should be inspected using *NeXpy* before proceeding (Figure 9C).

Note that in *mdx2* version 0.3.1, `mdx2.mask_peaks` does not account for systematic absences. This is a particular issue for space group I2_1_3, because the reflection condition *h* + *k* + *l* = 2*n* means that half of the Bragg peaks are absent. Although the mask generated above excludes twice as many spots as necessary, in this case relatively few pixels are excluded so the effect on the final map is minimal.

### 3.4. Background subtraction

Diffuse scattering experiments are different from Bragg data collection in that background must be measured carefully and subtracted (Pei et al., 2023). For insulin, background images were collected using 1 second exposures every 1 degree of rotation (compared with 0.1 second / 0.1 degree for the crystal). The background dataset was previously imported using *DIALS* in Section 3.1 to produce the file `background.expt`. First, import the dataset using `mdx2.import_data`

~~~
mdx2.import_data background.expt --outfile bkg_data.nxs --chunks 10 211 493
~~~

Inspect the images in `bkg_data.nxs` using *NeXpy*. The background pattern includes scattering from air, diffuse rings from the capillary surrounding the crystal, and a shadow of the pin (Figure 10A).

**Figure 10.**
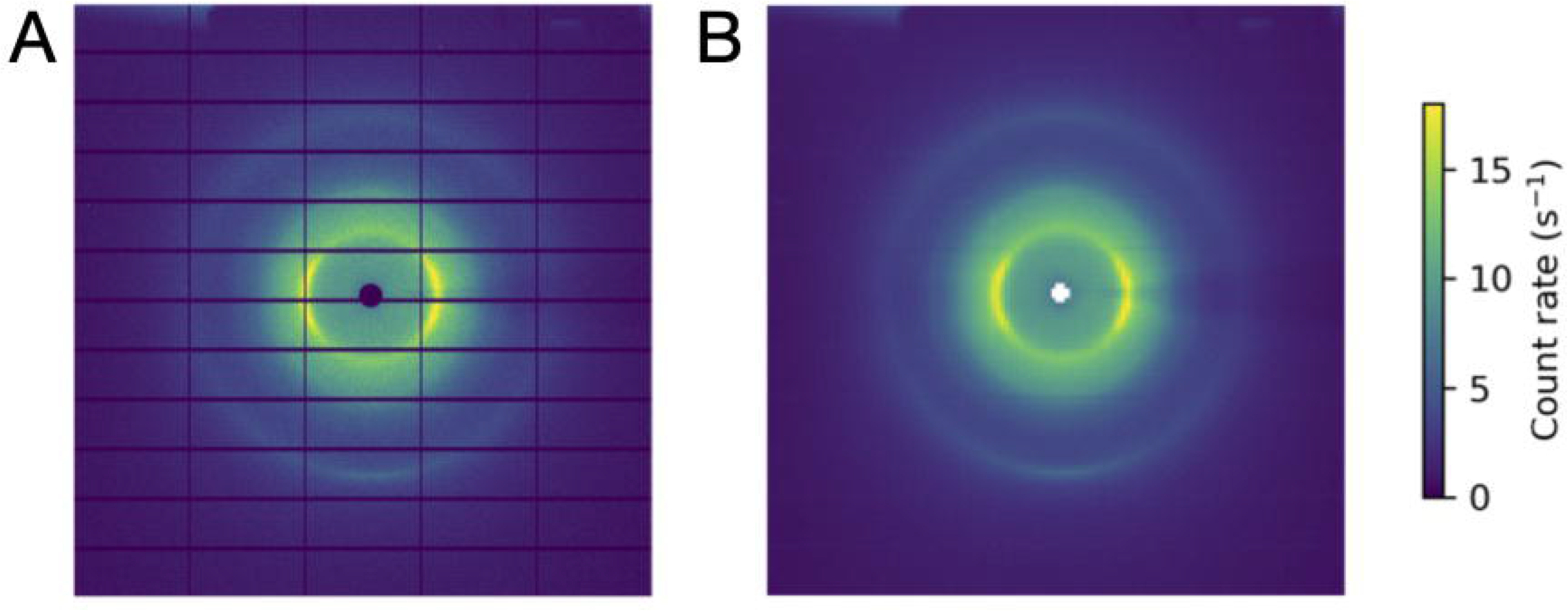
Image processing for background subtraction. (A) An example background image (1 degree oscillation, 1 second exposure) from the insulin rotation dataset. (B) The same image after binning to improve signal-to-noise. Background corrections are computed for each voxel in the reciprocal space map by interpolation of the binned image stack.

Because the background features vary gradually across the detector and with rotation angle, the noise can be reduced by smoothing. In *mdx2*, a downsampling (binning) method is used.

~~~
mdx2.bin_image_series bkg_data.nxs 10 20 20 --valid_range 0 200 --outfile
bkg_data_binned.nxs
~~~

The function has three mandatory arguments after the input file name to specify the bin size (here, 10 degrees by 20 pixels by 20 pixels). In addition, the optional argument `valid_range` is used to mask any pixel with counts outside the given interval. Here, a maximum of 200 counts was chosen to be 10 times the nominal background level of ∼20 counts per pixel (the background level is 10-fold higher in these images than in the crystal images due to the longer exposure time per frame). The threshold is used to reject broken pixels and stray diffraction if present (e.g., from tiny salt crystals), and is similar to `count_threshold` in `mdx2.find_peaks` described in Section 3.3.

The resulting binned image stack is saved as `bkg_data_binned.nxs` and can be viewed using *NeXpy* (Figure 10B). The binned images will be used after integration when applying geometric corrections according to Equation 15 (see Section 3.5).

### 3.5. Integration, Scaling and Merging

#### Integration

Integration refers to accumulating photon counts on a three-dimensional reciprocal space grid.

For the tutorial, a grid was chosen with 4 subdivisions in each direction, meaning voxels will be centered at Miller indices *h* = 1, 1.25, 1.5, 1.75, etc. This is by no means the optimal choice: the limit depends on the mosaicity, experimental geometry, and most importantly on how the diffuse scattering map will be used. However, when experimenting with different choices for the subdivisions, be aware that finer maps require more memory and disk space.

~~~
mdx2.integrate geometry.nxs data.nxs --mask mask.nxs --subdivide 4 4 4
~~~

For each voxel, the integrated photon counts and other data used in corrections are stored in a table in `integrated.nxs`. After integration, the pre-computed geometric and background corrections are applied (Equation 15):

~~~
mdx2.correct geometry.nxs integrated.nxs --background bkg_data_binned.nxs
~~~

The corrected intensities are saved as a table in `corrected.nxs`. The following Python code can be used to load the table as a *pandas* dataframe and preview the first several lines:

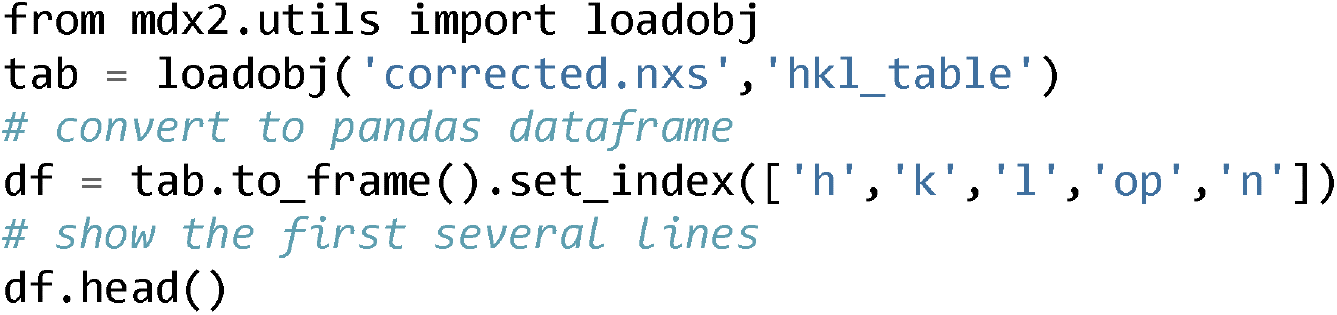

The columns contain the following information:

- h, k, l: Miller indices mapped to the *asymmetric unit*
- `op`: index of the symmetry operator that mapped the observation point to the asymmetric unit
- s: scattering vector magnitude (= 1/resolution)
- `intensity`, `intensity_error`: measured intensities and errors corrected using Equation 15.
- `rs_volume`: volume of reciprocal space recorded as a fraction of the reciprocal unit cell. Here, a voxel that is fully recorded would have `rs_volume` = 4^-3^ ∼ 0.0156.

#### Scaling

Finally, the corrected intensities can be scaled and merged. *Mdx2* implements the scaling algorithm from *mdx-lib* described in Section 2.3, however in *mdx2* version 0.3.1 only the per-image scale factor *b* is included in the model (Equation 21). Refine *b* as follows:

~~~
mdx2.scale corrected.nxs --smoothness 1
~~~

The model is refined using regularization to ensure that the correction factors vary gradually. The `smoothness` parameter sets the relative weight for minimizing smoothness vs. *χ*^2^. A value of 1 is a good starting point, but it can be varied up or down to observe the effect (e.g., by factors of 10). Additional optional parameters are available to further define the scaling model and refinement procedure (see `mdx2.scale --help`). By default, 5 cycles of alternating least-squares are performed and the scaling model is written to the file `scales.nxs`. To plot the refined scale factors (Figure 11), open the file in *NeXpy* and double-click `scaling_model` in the file tree.

**Figure 11.**
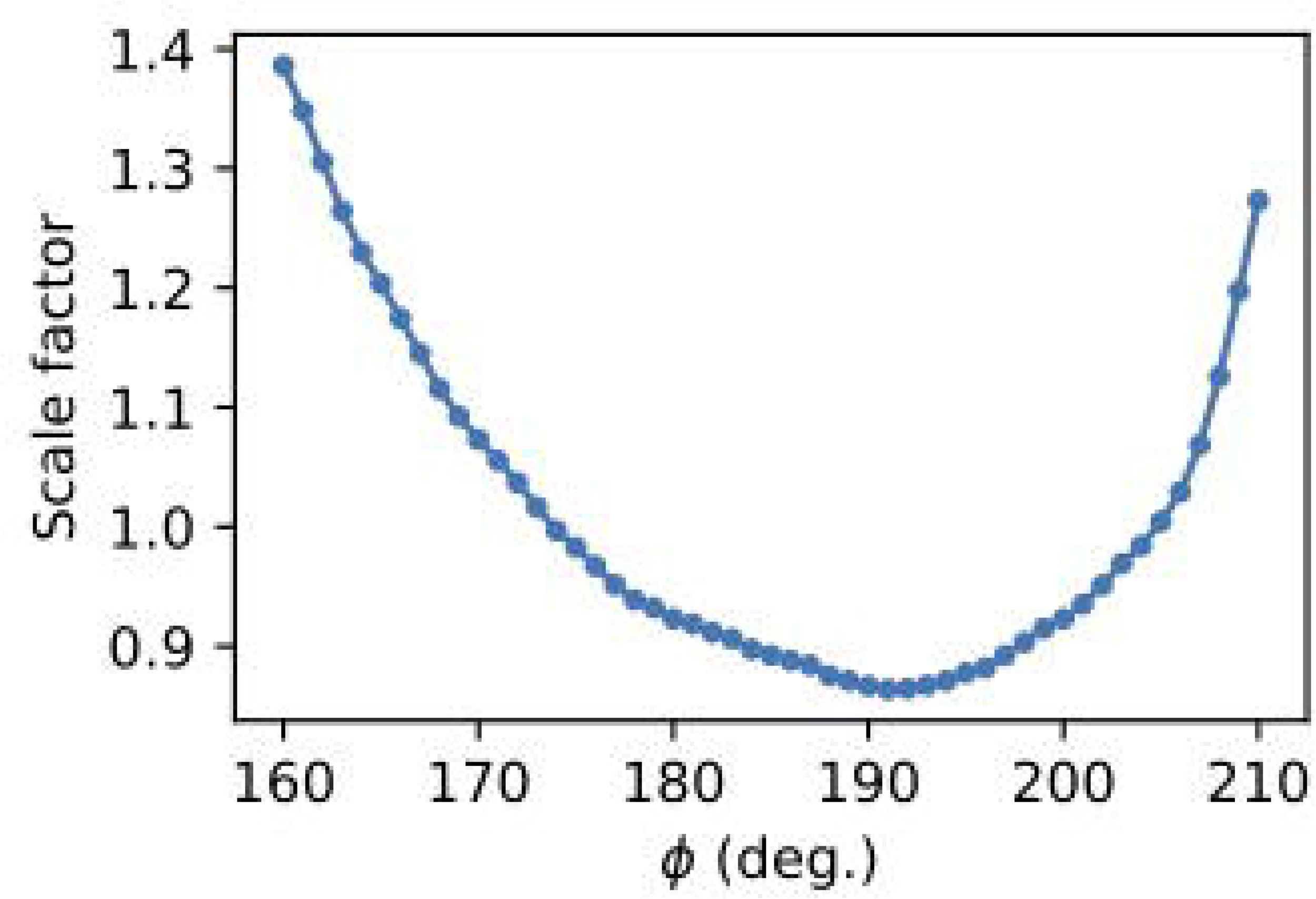
Scale factor per image in a rotation set after refinement using *mdx2*.

#### Merging

After scaling, the equivalent observations can be merged using Equations 19 and 20 as follows:

~~~
mdx2.merge corrected.nxs --scale scales.nxs
~~~

The file `merged.nxs` is produced with a table of intensities and error estimates.

### 3.6. Visualization

The processed diffuse scattering data from Section 3.5 is stored as a table of values covering the asymmetric unit of reciprocal space. In order to visualize the data, it is necessary to convert from a table to an array and symmetry-expand the data. The function *mdx2*.*map* can be used for generating slices and volumes. First, generate a slice as follows:

~~~
mdx2.map geometry.nxs merged.nxs --limits -50 50 -50 50 0 0 --outfile slice.nxs
~~~

The `--limits` argument defines the boundaries of the slice in fractional (Miller index) units. In this case, the slice is a central section (*hk0*) with *h* from -50 to 50 (fractional units), *κ* from -50 to 50 (fractional units), and *l* = 0. With the 4 × 4 × 4 sampling used, the slice will have 201 × 201 voxels.

Open the file `slice.nxs` in *NeXpy* and double-click `intensity` in the file tree to display a slice through the reciprocal space map. Adjust the sliders in the signal tab to make the background scattering variations clear (as in Figure 12).

**Figure 12.**
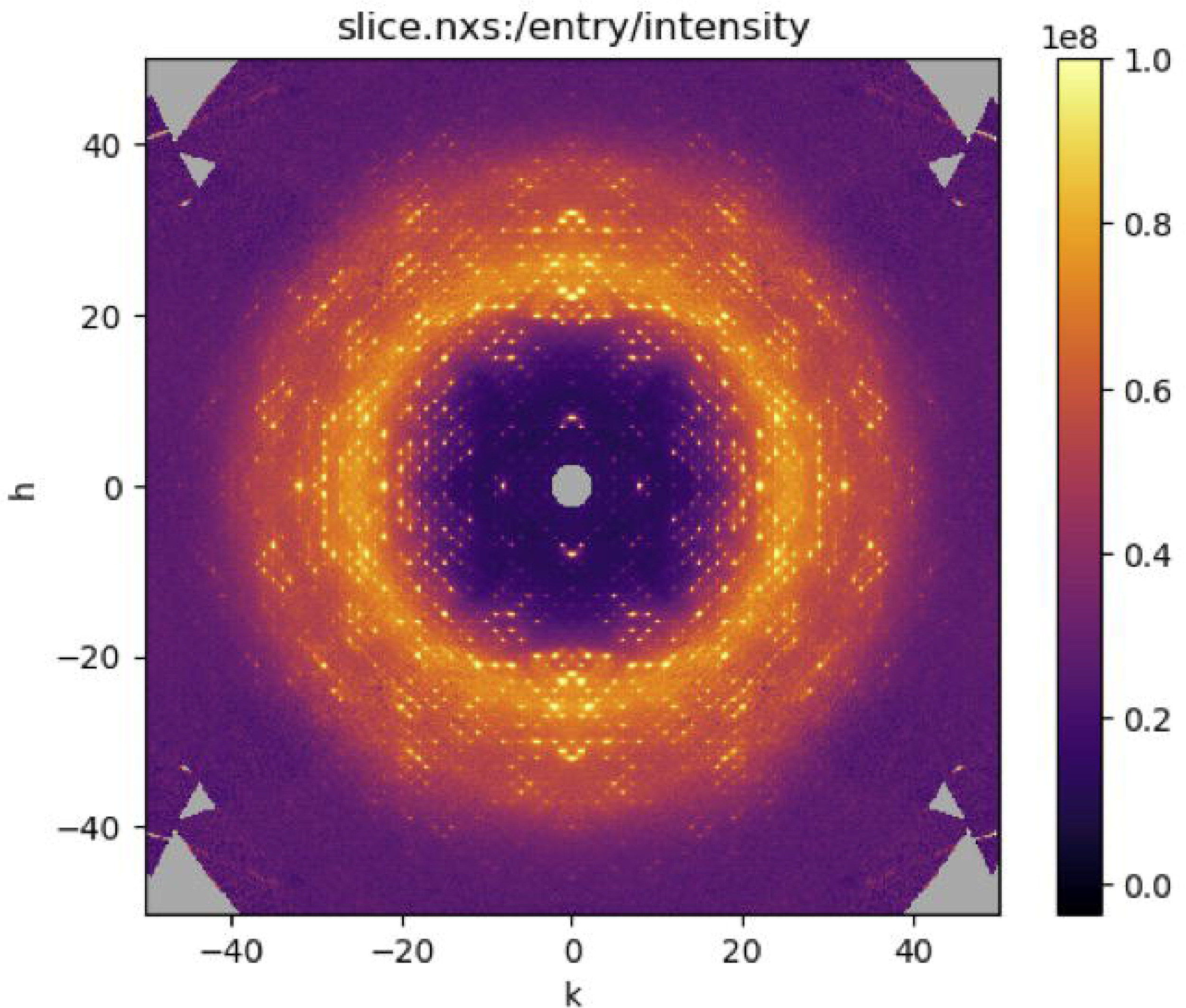
Slice through the reciprocal space map of insulin visualized using *NeXpy*.

A similar command can be used to generate reciprocal space volumes:

~~~
mdx2.map geometry.nxs merged.nxs --limits 0 50 0 50 0 50 --outfile map.nxs
~~~

To conserve computing resources, we have restricted the map to the positive octant of reciprocal space (*h* ≥ 0, *κ* ≥ 0, *l* ≥ 0), which is the same as the other seven by symmetry. Open `map.nxs` using *NeXpy* and double-click `intensity` in the file tree. Scroll through *z*-slices. It helps to unselect “autoscale” in the “z” tab so that intensity scale is fixed, and then adjust it in the “signal” tab.

### 3.7. Map Statistics

In this section, we use *mdx2* with *pandas* to calculate statistical properties of the diffuse map. *Pandas* is a Python library commonly used in data science to manipulate tabular data. It will be used here without explanation; excellent tutorials are available elsewhere (VanderPlas, 2016).

In a Python session or Jupyter notebook, first import the needed libraries and define convenient functions as follows:

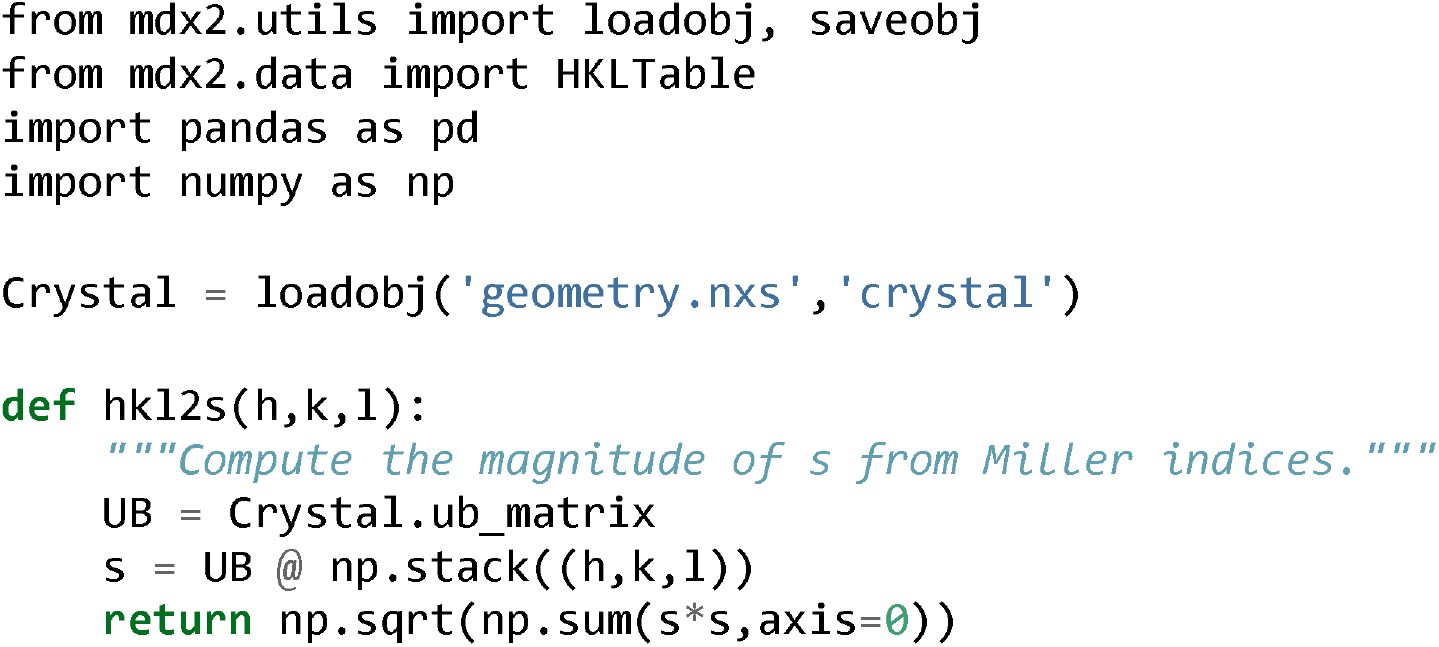

#### Intensity vs. resolution

The following Python program calculates the mean and standard deviation of intensity in shells of constant resolution (Figure 13A):

**Figure 13.**
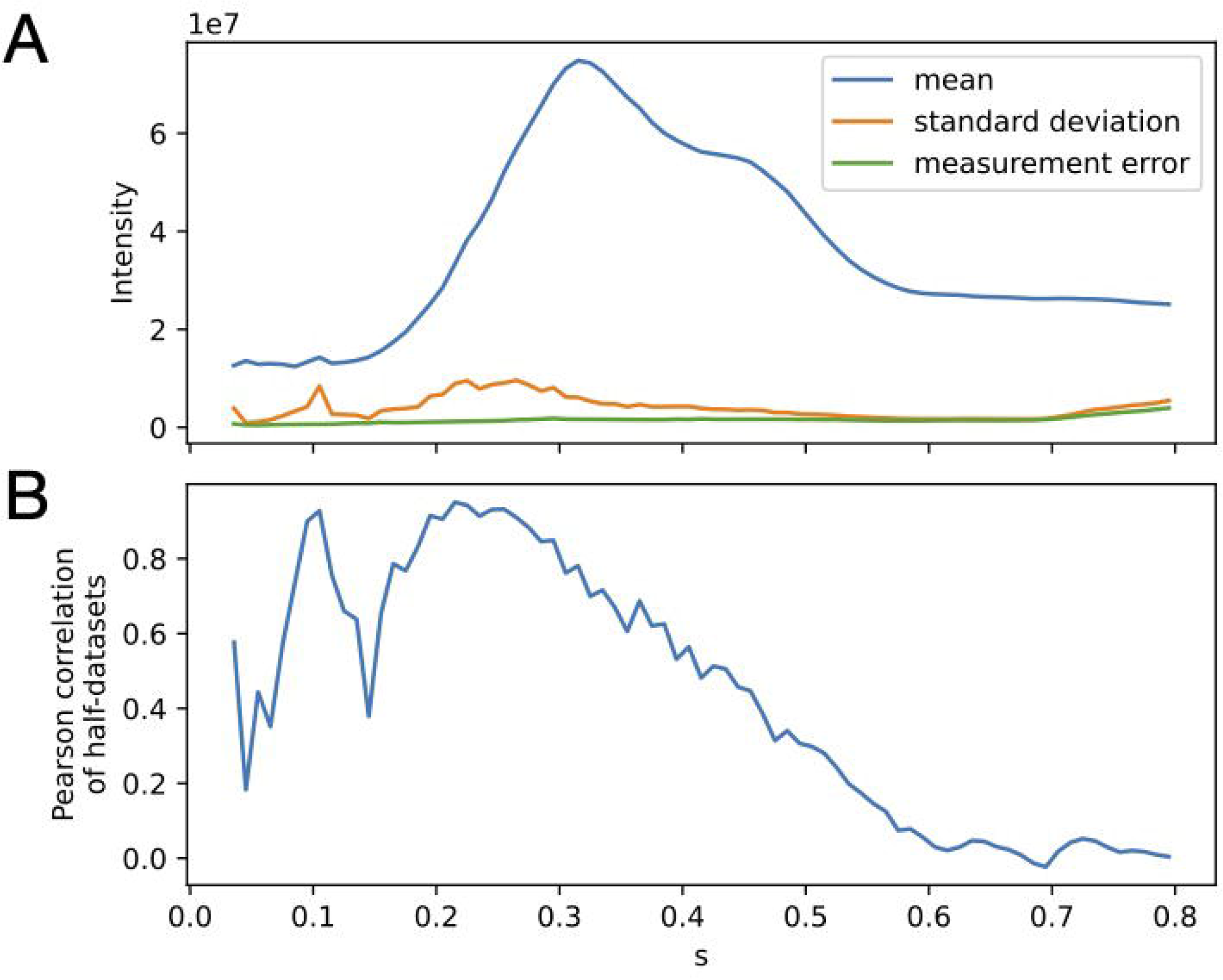
Intensity statistics for the insulin diffuse scattering dataset after scaling. (A) The mean (blue) and standard deviation (orange) of intensity were computed within each resolution shell for the merged reciprocpal space map. The variations in the diffuse pattern are much smaller than the isotropic component. However, they are larger than the noise in the data (average measurement error, propagated from Poisson statistics), shown in green. (B) The dataset was split into equivalent halves and merged separately, and the Pearson correlation was computed between these half datasets as a function of resolution (1/s). By this measure, the variations in the diffuse pattern contain signficant signal up to ∼2 Å resolution. The dip at low resolution is related to the strength of the halo features relative to the background in this dataset.

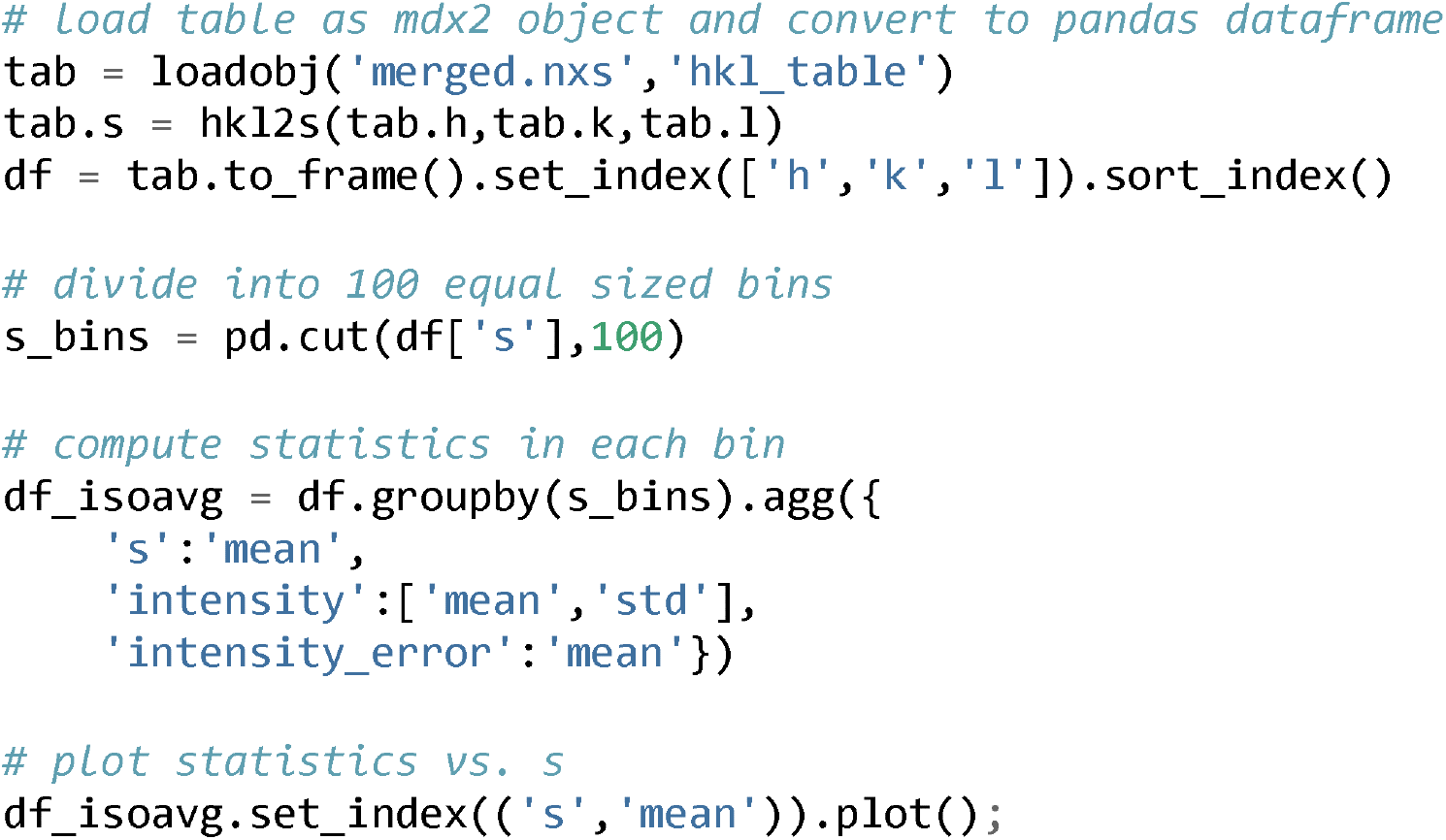

#### Correlation of half-datasets

To quantify the precision of the final diffuse map, the dataset can be split into equivalent halves and the correlation coefficient computed between them in shells of constant resolution (see Chapter 1 (Pei et al., 2023)). *Mdx-lib* has built-in tools for splitting unmerged datasets randomly with similar statistical weight. This method is not yet available in *mdx2* version 0.3.1. Here, we use *pandas* to split the data by assigning asymmetric units to each half (splitting on the op column of the data table). The following python script reads the unmerged data from `corrected.nxs` and saves two files: `split_1.nxs` and `split_2.nxs`.

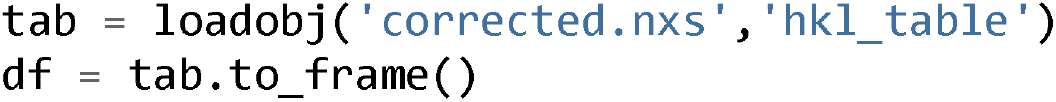
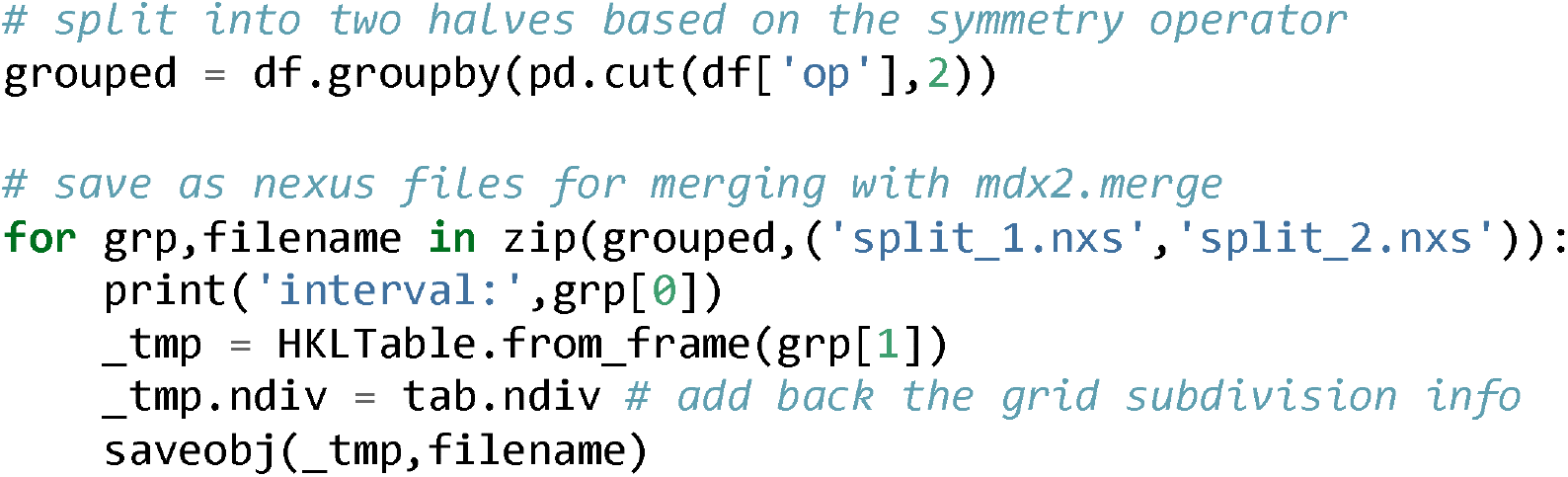

Next, scale and merge each half dataset using *mdx2* on the command line:

~~~
mdx2.merge split_1.nxs --scale scales.nxs --outfile merged_1.nxs
mdx2.merge split_2.nxs --scale scales.nxs --outfile merged_2.nxs
~~~

Finally, load the merged datasets as *pandas* dataframes and compute correlation coefficients in shells of constant resolution (Figure 13B):

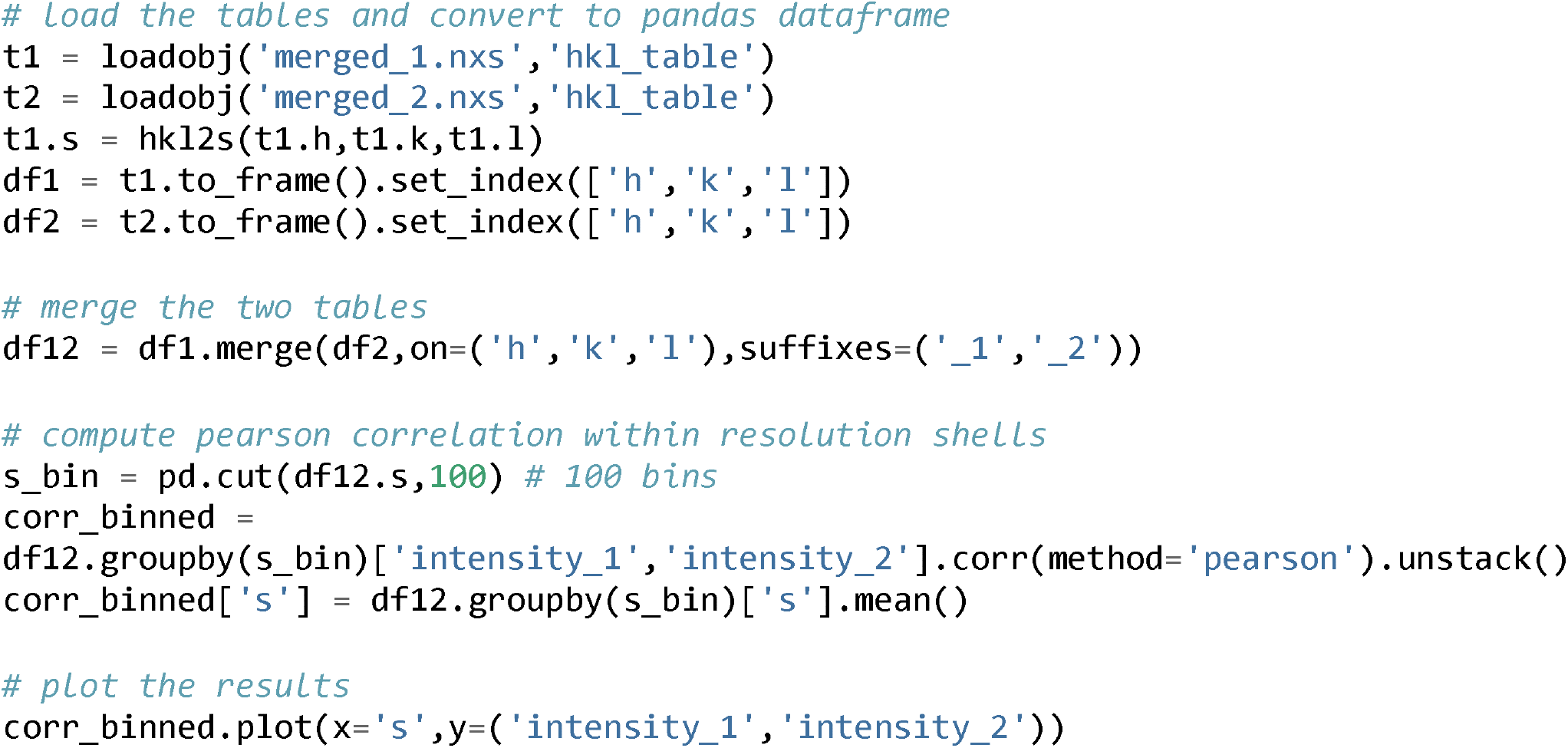

## 4. Conclusions

This chapter has introduced a modern approach to macromolecular diffuse scattering based on low dose, high redundancy data collection and reciprocal space mapping. The promise of diffuse scattering for studying correlated motion was made clear by Caspar and colleagues 35 years ago, however it has only recently become feasible to acquire the accurate and detailed data needed to account for all of the motions occuring in a protein crystal. Much of the recent work on diffuse scattering has focused on data processing, and we expect a continued focus on methods development in the near future. However, the field has reached a turning point. As outlined in Chapter 1 (Pei et al., 2023), now that the physical basis for diffuse scattering from protein crystals is understood, the demands on sample quality and data processing have been clarified. Although methods will continue to evolve, we expect increased standardization of software and workflows going forward. Here, we have made the first steps toward standardization in the form of *mdx2*. Furthermore, by introducing user-friendly software tools and rigorous tutorials, we have reduced the burden of expertise required to enter the field of diffuse scattering. With these advances, the goal of moving the field beyond proof-of-concept experiments to biological applications is within reach.

## Acknowledgments

The *mdx2* software and tutorial in Section 3 are based on materials developed for the Data Reduction Workshop at the 2022 Erice School on Diffuse Scattering. We are grateful to the organizers and participants for providing both the motivation to develop *mdx2* and valuable feedback. We thank Kevin Dalton and members of the *DIALS* development team for answering questions and helping us get started with *dxtbx*. The insulin dataset was collected at the Center for High-Energy X-ray Sciences (CHEXS), which is supported by the at National Science Foundation (BIO, ENG and MPS Directorates) under award DMR-1829070, and the Macromolecular Diffraction at CHESS (MacCHESS) facility, which is supported by award P30GM124166 from the National Institute of General Medical Sciences, National Institutes of Health, and by New York State’s Empire State Development Corporation (NYSTAR). This work was supported by National Institutes of Health grant GM124847 (to N.A.).

## Appendix A

### A.1. Obtaining and running the tutorial in Jupyter notebooks

The tutorials in Section 3 are available on the *mdx2* GitHub page as separate Jupyter notebooks (https://github.com/ando-lab/mdx2). The notebooks can be downloaded individually or obtained by cloning the *mdx2* repository.

The code is designed to run on a personal computer with ∼20 Gb of free disk storage and at least 4 Gb of RAM. A unix-like operating system is assumed (Linux, OSX, or Windows Subsystem for Linux).

### A.2. Downloading the tutorial dataset

The dataset from insulin is available on Zenodo (https://dx.doi.org/10.5281/zenodo.6536805). First, download insulin_2_1.tar and extract the tar archive. The file will expand to a directory called `images` with subfolders `insulin_2_1`, `insulin_2_bkg`, and `metrology`. Place `images` in the same directory as the Jupyter notebooks.

### A.3 Setting up the python environment

Install *miniconda3* with *python* version *3*.*10*. Installers and instructions are available on the web: https://docs.conda.io/en/latest/miniconda.html. To prevent miniconda from interfering with existing conda installations, it should be prevented from modifying the users’ bash scripts or running “conda init” (-b flag). We also recommend choosing a non-default install location; `∼/miniconda-mdx2` is assumed in the following examples, but can be modified if needed.

Example (MacOS x64):

~~~
wget https://repo.anaconda.com/miniconda/Miniconda3-py310_22.11.1-1-MacOSX-
x86_64.sh -O ∼/miniconda-installer.sh
bash ∼/miniconda-installer.sh -b -p $HOME/miniconda-mdx2
~~~

Next, activate the *mdx2* conda environment and install *mdx2*’s dependencies

~~~
source ∼/miniconda-mdx2/bin/activate
conda install -c conda-forge dxtbx nexusformat pandas numexpr
~~~

If the conda environment is de-activated (by typing `conda deactivate` or starting a new terminal session), it can be re-activated by typing source ∼/miniconda-mdx2/bin/activate. For convenience, the following line can be added to the shell startup script:

Example (`∼/.bash_profile`)

~~~
alias activate_mdx2=“source ∼/miniconda-mdx2/bin/activate”
~~~

Then, the mdx2 environment can be activated by typing `activate_mdx2`.

### A.4. Installing mdx2

First, activate the *mdx2* conda environment (see above). Then install *mdx2* from the GitHub repository using pip

~~~
pip install git+https://github.com/ando-lab/mdx2.git@v0.3.1
~~~

The tag @v0.3.1 at the end of the repository address specifies version 0.3.1 for consistency with this tutorial. If this tag is omitted, the most up-to-date version will be installed (see Version notes at https://github.com/ando-lab/mdx2).

The *mdx2* tools should now be available at the command line. Check the version as follows:

~~~
mdx2.version
~~~

The version number 0.3.1 should be displayed.

### A.5. Installing DIALS, NeXpy, and JupyterLab

First, activate the *mdx2* conda environment (see above). Then install the programs using conda

~~~
conda install -c conda-forge dials nexpy jupyterlab
~~~

With the *mdx2* conda environment active, *nexpy* can be launched from the command line by typing `nexpy`. Similarly, Jupyter Lab can be launched by typing `jupyter lab`. The *DIALS* command-line tools are also available. For instance `dials.version` prints the version information and install location.

